# Genetic Analysis of Activin/Inhibin β Subunits in Zebrafish Development and Reproduction

**DOI:** 10.1101/2022.09.23.509125

**Authors:** Cheng Zhao, Yue Zhai, Ruijing Geng, Kun Wu, Weiyi Song, Nana Ai, Wei Ge

## Abstract

Activin and inhibin are both dimeric proteins sharing the same β subunits that belong to the TGF-β superfamily. They are well known for stimulating and inhibiting pituitary FSH secretion, respectively, in mammals. In addition, activin also acts as a mesoderm-inducing factor in frogs. However, their functions in development and reproduction of other species are poorly defined. In this study, we disrupted all three activin/inhibin β subunits (βAa, *inhbaa*; βAb, *inhbab*; and βB, *inhbb*) in zebrafish using CRISPR/Cas9. The loss of βAa/b but not βB led to a high mortality rate in the post-hatching stage. Surprisingly, the expression of *fshb* but not *lhb* in the pituitary increased in the female βA mutant together with aromatase (*cyp19a1a*) in the ovary. The single mutant of βAa/b showed normal folliculogenesis in young females; however, their double mutant (*inhbaa−/−*;*inhbab−/−*) showed delayed follicle activation, granulosa cell hypertrophy, stromal cell accumulation and tissue fibrosis. The ovary of *inhbaa−/−* deteriorated progressively after 180 dpf with reduced fecundity and the folliculogenesis ceased completely around 540 dpf. In addition, tumor- or cyst-like tissues started to appear in the *inhbaa−/−* ovary after about one year. In contrast to females, activin βAa/b mutant males showed normal spermatogenesis and fertility. As for activin βB subunit, the *inhbb−/−* mutant exhibited normal folliculogenesis, spermatogenesis and fertility in both sexes; however, the fecundity of mutant females decreased dramatically at 270 dpf with accumulation of early follicles. In summary, the activin-inhibin system plays an indispensable role in fish reproduction, in particular folliculogenesis and ovarian homeostasis.

## Introduction

Activin and inhibin are both dimeric proteins containing β subunits that belong to the transforming growth factor β (TGF-β) superfamily. Activins are homodimers of βsubunits (βAβA, βBβB and βAβB) whereas inhibins are heterodimers with one β and one unique α subunit (αβA and αβB) [1, 2]. These molecules were discovered in the ovarian follicular fluid for their specific stimulation and inhibition of pituitary follicle-stimulating hormone (FSH) secretion, respectively [3, 4]. Despite sharing the same β subunits, activin and inhibin are functionally antagonistic in nearly all target cells [2]. In addition to regulating FSH secretion and therefore reproduction in mammals, activin has also been found to serve as a potent mesoderm-inducing factor (MIF) during embryonic development in the *Xenopus* [5, 6]; however, this role has not been supported by genetic studies in mammals [7–9], making it a highly controversial issue in developmental biology. It would be interesting to address this question in other organisms such as zebrafish, which is one of the top vertebrate models for both genetics and developmental biology.

Activin acts by binding to specific receptors, activin type I (ActRI) and type II (ActRII) receptors, which trigger the intracellular signal transduction pathways involving primarily Smad proteins. Activin first binds to a type II receptor followed by recruiting and activating a type I receptor. The type I receptor in turn induces phosphorylation of Smad2 and/or Smad3, which then recruits a common partner Smad4 to form a complex for translocation into the nucleus to regulate expression of the target genes [10, 11]. As a natural antagonist of activin, inhibin works by competing with activin for binding the same type II receptors but without activating the type I receptors [12, 13].

The discovery of activin-inhibin system has been considered one of the most important breakthroughs in reproductive endocrinology in 1980s [1]. In addition to regulating pituitary FSH biosynthesis, activin also acts as a local factor in the gonads to regulate gametogenesis and steroidogenesis in autocrine and/or paracrine manners. In postnatal mice, activin promotes primordial follicle formation by stimulating germline cyst breakdown or follicle assembly [14]. In mature females, activin promotes differentiation of primordial follicles into antral follicles and stimulates granulosa cell proliferation [15]. In cultured ovarian granulosa or granulosa-lutein cells, activin increases expression of FSH receptor [16, 17] but decreases that of LH receptor [17]. Activin is also involved in regulating steroidogenesis in follicles. It stimulates expression of aromatase and estrogen receptors [16, 17] but decreases expression of steroidogenic acute regulatory protein (StAR) and production of progesterone in the granulosa cells [17] as well as androgen production in the theca cells [18]. In the testis, activin induces Sertoli cell proliferation in culture alone or in combination with FSH [19, 20] and regulates steroid production [21].

The functional importance of activin/inhibin β subunit genes (hereafter referred to as activin β subunits) in mammals has also been studied by genetic approaches. Knockout of activin βA subunit (*Inhba*) in mice showed defects in craniofacial development and failed growth of whiskers with lower incisors and cleft palate at birth, and all mutant individuals died within 24 h after birth, making it impossible to study its functions in reproduction [8]. Disruption of activin βB subunit (*Inhbb*) resulted in defective eyelid closure and prolonged gestation; however, the mutant could survive with normal growth and gonadal development [8, 22]. Conditional knockout of *Inhba* in ovarian granulosa cells did not cause mortality, but the mutant was subfertile, suggesting important roles for gonadal βA in ovarian function. Double mutation of *Inhba* and *Inhbb* in the ovary (conditional knockout of *Inhba* in the ovary in the background of global null knockout of *Inhbb*) resulted in complete infertility [23]. These results suggest that activin βA and βB are both required for normal ovarian folliculogenesis, with βA playing a more prominent role. In the testis, activin is primarily expressed in the Sertoli cells, but its signal could also be detected in a variety of other cell types, including Leydig and spermatogenic cells as well as other somatic cells in various mammalian species [24]. During embryonic development, activin is mainly produced by fetal Leydig cells and acts on Sertoli cells to promote their proliferation [25, 26], and specific disruption of *Inhba* in Leydig cells resulted in decreased proliferation of Sertoli cells [26].

In teleosts, activin βA and βB subunits were first cloned in goldfish (*Carassius auratus*), and they are widely expressed in both gonadal and non-gonadal tissues [27, 28]. In rainbow trout (*Oncorhynchus mykiss*), the βA and βB are expressed in the theca layer in the ovary and interstitial cells in the testis [29]. In zebrafish, there exist three β subunits (βAa, *inhbaa*; βAb, *inhbab*; and βB, *inhbb*), and they are exclusively expressed in the somatic follicle layer surrounding the oocytes in the ovary whereas activin receptors (type II and I) and downstream signaling molecules Smads are abundantly expressed in the oocytes, suggesting a potential paracrine pathway for follicle cell-to-oocyte signaling [30, 31]. During folliculogenesis, activin subunits exhibit dynamic expression patterns. The expression of *inhbaa* increased steadily during follicle growth phase, starting from follicle activation, *i.e.*, the transition from primary growth (PG) to secondary growth (SG; starting from previtellogenic stage, PV). The expression reached the maximum level at mid-vitellogenic (MV) stage followed by a decrease at the full-grown (FG) stage [32, 33]. Interestingly, *inhbab* (a paralogue of βA) exhibited a different expression pattern from *inhbaa*. It increased steadily during folliculogenesis and reached the maximum level at the FG stage [34]. The expression of *inhbb* remained largely unchanged during follicle development; however, it showed a sharp increase in the ovary prior to spawning [32]. Despite these studies, the functions of activin in fish reproduction remain poorly understood. Activin stimulates pituitary FSHβ (*fshb*) but suppresses LHβ (*lhb*) expression in several fish species including zebrafish [35], goldfish [36] and eel [37], in contrast to its stimulation of FSH only in mammals [1]. Activin also works in the gonads as a local paracrine factor. In males, activin stimulates proliferation of spermatogonia in the testis of Japanese eel (*Anguilla japonica*) [38]. In females, activin promotes the acquisition of oocyte maturational competence and stimulates final maturation in zebrafish [39, 40] and killifish [41], and it suppresses both basal and hCG-stimulated secretion of testosterone in early vitellogenic but not FG follicles in goldfish [42]. As a local ovarian factor expressed in the somatic follicle cells, activin is likely regulated by both endocrine and paracrine factors that act or interact on the follicle cells. Activin may mediate actions of pituitary gonadotropins as its expression was significantly up-regulated by gonadotropin(s) in zebrafish [43]. In addition, activin subunits are also subject to regulation by local factors. Epidermal growth factor (EGF) from the oocyte significantly stimulated expression of all three subunits in cultured zebrafish follicle cells [44]. Although all lines of evidence suggest roles for activins in fish reproduction, their exact activities and functional importance remain largely unknown due to the lack of genetic studies in fish models.

Using CRISPR/Cas9 method, we carried out a comprehensive genetic analysis in the present study to investigate the functions of activin β subunits in zebrafish focusing on female reproduction. All three β subunit genes (*inhbaa, inhbab* and *inhbb*) were deleted followed by phenotype analysis on single, double, and triple knockouts of these genes. The results suggest important roles for activin β subunits in controlling pituitary gonadotropin expression, follicle activation and ovarian homeostasis.

## Materials and Methods

### Animals and maintenance

The wild type (WT) zebrafish of AB strain was used in the present study to generate mutant lines. Adult zebrafish were maintained in the ZebTEC Multilinking Rack zebrafish system (Tecniplast, Buguggiate, Italy) at 28°C and pH 7.5 with a lighting scheme of 14-h light (8:00 am to 10:00 pm) and 10-h dark, and they were fed with Otohime fish diet (Marubeni Nisshin Feed, Tokyo, Japan) delivered by the Tritone Automatic Feeder system (Tecniplast). Zebrafish fry were fed with paramecium from 5 to 10 days post-fertilization (dpf) and artemia from 10 to 15-20 dpf in nursery tanks until transfer into the ZebTEC aquarium system. All animal experiments were conducted in accordance with the guidelines and approval of the Animal Ethics Panel of the University of Macau.

### Establishment of mutant zebrafish lines for activin β subunits

The genomic and coding sequences of zebrafish *inhbaa*, *inhbab* and *inhbb* were obtained from the Ensembl (asia.ensembl.org/*Danio_rerio*). Single mutants of *inhbaa*, *inhbab* and *inhbb* were generated by CRISPR/Cas9-induced mutagenesis. Briefly, the CRISPR target sites were designed using the online tool (ZiFiT Targeter Version 4.2; http://zifit.partners.org/ZiFiT/). The primer pairs for CRISPR/Cas9 knockout are shown in Table S1. The Cas9 mRNA and single-guide RNAs (sgRNAs) were generated with the mMACHINE T7 and mMACHINE SP6 kits (Invitrogen, Carlsbad, CA). The Cas9 mRNA (300 ng/µL) and sgRNA (50 ng/µL) were co-microinjected into one- or two-cell-stage embryos (4.6 nL/embryo) using the Drummond Nanoject injector (Drummond Scientific, Broomall, PA). The embryos were sampled after 24 h for high-resolution melt analysis (HRMA) and heteroduplex motility assay (HMA) to check mutagenesis. The F0 founders carrying mosaic mutations were raised to the adult stage and outcrossed with WT fish to generate heterozygous F1 offspring as we previously described [45].

### Genotyping by HRMA and HMA

Briefly, the genomic DNA from an embryo or a piece of caudal fin was extracted by the NaOH method [46, 47] and used as the template for HRMA as described [47, 48]. Primers flanking the target site were used to amplify the target region of the gene in a 10 µL polymerase chain reaction (PCR) containing: 5 µL 2 × SuperMix (Bio-Rad, Hercules, CA), 1 µL genomic DNA, and 0.2 µM forward and reverse primers (Table S1). The HRMA conditions were 95°C for 3 min; 40 cycles of 95°C for 15 sec, 60°C for 15 sec, and 72°C for 20 sec, followed by a final melting curve analysis from 70°C to 95°C with 0.2°C increment for each step using the CFX96 Real-Time PCR System and the Precision Melt Analysis software (Bio-Rad). For HMA analysis, the HRMA product (7 µL) was electrophoresed on 20% polyacrylamide gel at 120 V for 5 h, stained with GelRed (Biotium, Hayward, CA) and visualized on Gel Doc System (Bio-Rad) [48, 49].

### Survival analysis

According to previous studies on activins in mice and *Xenopus*, we speculated that the loss of activin β subunits would affect embryonic and larval development, and therefore the survival of zebrafish. To test this, we examined the ratios of different genotypes in the offspring from incrossing between heterozygotes (+/−). We also characterized the time window when the lethality occurred for single and βA double mutants by analyzing the ratio of each genotype from 1 to 22 dpf. To obtain single mutants like *inhbaa−/−*, we crossed the heterozygotes [*inhbaa*+/− (♀) x *inhbaa*+/− (♂)]. To obtain double mutants like *inhbaa*−/−*;inhbab*−/−, the crossing was made as follows: *inhbaa*+/−*;inhbab*−/− (♀) x *inhbaa*−/−*;inhbab*+/− (♂). To obtain βA and βB triple mutant (*inhbaa*−/−*;inhbab*−/−*;inhbb*−/−), the crossing was made between *inhbaa*+/−*;inhbab*−/−*;inhbb*+/− (♀) and *inhbaa*−/−*;inhbab*+/−*;inhbb*−/− (♂). The larval fish were sampled at different time points for DNA extraction and genotype analysis.

For survival analysis of activin βA mutant after 50 dpf, we bred the fish as follows: *inhbaa*+/−*;inhbab*−/− (♀) x *inhbaa*−/−*;inhbab*+/− (♂), and the offspring were genotyped at 45 dpf to identify four genotypes (*inhbaa*+/−*;inhbab*+/−, *inhbaa*−/−*;inhbab*+/−, *inhbaa*+/−*;inhbab*−/− and *inhbaa*−/−*;inhbab*−/−), which were then raised in separate tanks under the same condition (46-54 fish/tank). The number of fish in each tank was monitored every five days from 50 to 165 dpf to estimate the survival rates of the βA mutants. To obtain enough homozygous double mutants of the βA subunits for analysis, we also bred the double mutants [*inhbaa*−/−*;inhbab*−/− (♀) x *inhbaa*−/−*;inhbab*−/− (♂)] in the iSpawner (Tecniplast) to increase breeding success to overcome the low fecundity and high mortality of the double mutant (*inhbaa*−/−*;inhbab*−/−).

### Separation of oocyte and follicle layer

To examine spatial distribution of gene expression in the two cell types of follicles (oocyte and follicle layer), we separated the somatic follicle layer from the oocyte of FG stage according to our previous reports [31, 50]. Briefly, the full-grown immature follicles were isolated and incubated in the Ca^2+^/Mg^2+^-free Cortland medium for 30 min at −20℃ to loosen the attachment between the oocyte and follicle layer. While the follicle was held gently by a blunt-end forceps, the follicle layer was carefully peeled off as a whole from the oocyte with a sharp forceps without breaking the oocyte. The isolated follicle layers and denuded oocytes from five follicles were pooled together for RNA extraction in TRIzol (Invitrogen, Carlsbad, CA).

### Sampling and histological analysis

Briefly, the fish were anaesthetized with MS-222 (Sigma-Aldrich, St. Louis, MO) and placed on a Petri dish cover to measure standard body length (BL) and body weight (BW). The entire fish or dissected gonad was immediately fixed in Bouin’s solution for at least 24 h at room temperature. The fixed samples were dehydrated using the ASP6025S Automatic Vacuum Tissue Processor (Leica, Wetzlar, Germany). The samples were imbedded in paraffin followed by serial sectioning at 5 µM thickness using the Leica microtome (Leica). The sections were deparaffinized, hydrated, stained with hematoxylin and eosin (H&E) and mounted with Canada balsam (Sigma-Aldrich). The sections were examined on the Nikon ECLIPSE Ni-U microscope with Digit Sight DS-Fi2 digital camera (Nikon, Tokyo, Japan).

To determine follicle composition in the ovary, we performed serial longitudinal sectioning on the whole fish at 5 µm. The largest ovarian section of each fish was chosen for follicle quantification by NIS-Elements BR software (Nikon). Follicle staging was based on both morphological features (cortical alveoli and yolk granules) and oocyte diameter according to our previous studies [32, 51]. Briefly, the process of folliculogenesis was divided into six stages: primary growth (PG, <150 µm), previtellogenic (PV, ∼250 µm), early vitellogenic (EV, ∼350 µm), mid-vitellogenic (MV, ∼450 µm), late vitellogenic (LV, ∼550 µm) and full-grown (FG, >650 µm) stages.

### RNA isolation and real-time qPCR

Total RNA was isolated from the follicle layers, denuded oocytes and different tissues using TRIzol (Invitrogen) and quantified based on the absorbance at 260 nm using the NanoDrop 2000 (Thermo Scientific, Waltham, MA). Reverse transcription (RT) was performed on 3 µg total RNA using MMLV reverse transcriptase (Invitrogen) according to the manufacturer’s instructions. The cDNA product was used as the template for PCR amplification. Semi-quantitative RT-PCR and agarose gel electrophoresis were used to localize the expression of the activin-inhibin-follistatin system in the follicle. The amplification was performed in a total volume of 15 μL containing 5 μL of 1:15 diluted RT products and standard PCR reagents. The cycle numbers of RT-PCR were optimized as previously reported [31]: 32 for *inhbaa*, *inhbab*, *gdf9* and *lhcgr*; 35 for *inhbb*, *fsta* and *fstb*; and 27 for *inha* and *ef1a*. Quantitative real-time PCR (qPCR) was used to determine expression levels of target genes in the pituitary, ovary and testis. The PCR reactions were performed on the CFX96 Real-Time PCR System (Bio-Rad) in a volume of 10 μL containing 5 μL 2× SuperMix, 0.2 μM each primer, and 4.8 µL cDNA template. Each sample was normalized to the expression level of housekeeping gene *ef1a* and the value was expressed as the fold change over the control group. The primers used in RT-PCR and qPCR are listed in Table S1.

### Fluorescent in situ hybridization

The zebrafish were sacrificed by decapitation after cold shock in ice water. The heads of zebrafish were fixed overnight at room temperature in 4% paraformaldehyde in phosphate-buffered saline (PBS), dehydrated in serial ethanol and embedded in paraffin. The head sections through the pituitary (5 μm) were mounted on slides. The PCR products of target genes were purified and used as templates for probe preparation using DIG or fluorescein RNA labeling kits (Roche, Basel, Switzerland). The pituitary sections were hybridized with the antisense RNA probes for *fshb* and *lhb* as we previously reported [52]. The TSA Plus Fluorescein/Cy5/TMR system (PerkinElmer, Waltham, MA) was used to detect the fluorescent signals.

### TUNEL staining for apoptosis

The gonads were dissected and immediately fixed in 4% paraformaldehyde in PBS for 24 h at room temperature followed by dehydration and paraffin embedding. The tissues were sectioned at 5 µm thickness followed by deparaffinization and rehydration. Then the sections were incubated with proteinase K in PBS (20 µg/mL) at 37°C for 20 min. TUNEL staining was performed with the In Situ Cell Death Detection Kit (fluorescein) (Roche) in a humidified chamber at 37 °C for 1 h.

### Masson trichrome staining and sirius red staining

To evaluate the fibrosis of the ovarian tissues, the ovaries were collected from different genotypes of zebrafish and fixed in 4% paraformaldehyde for 48 h. Following paraffin embedding and sectioning (5 μm), the tissues were deparaffinized, rehydrated and stained with Masson trichrome and Sirius red as reported [53]. For Masson trichrome staining, the sections were stained sequentially with Weigert’s working hematoxylin, Biebrich scarlet-acid fuchsin, phosphomolybdic-phosphotungstic acid, aniline blue and acetic acid solutions for 5 min each (Phygene Life Sciences, Fuzhou, China). The collagen fibers are stained blue with Masson trichrome staining. For sirius red staining, the sections were stained in picro-sirius red (Phygene Life Sciences, Fuzhou, China) for 1 h followed by washing in acetic acid solution and absolute alcohol. Collagen fibers are stained red with sirius red staining.

### Determination of proinflammatory factors

To assess inflammation in ovarian tissues, we determined the levels of tumor necrosis factor-α (TNF-α) and interleukin 6 (IL-6) in the ovary by ELISA (Renjie Biotechnology, Shanghai, China) according to manufacturer’s instructions and a previous study [54]. The antibodies used for the ELISA were generated specifically with full-length recombinant zebrafish TNF-α and partial sequence of recombinant zebrafish IL-6 (amino acids 87-231). Briefly, 30 µg ovarian sample from each fish (4 fish in total for each group) was homogenized in 300 μL ice-cold saline buffer, centrifuged for 10 min (3000 × g) and the supernatants (50 µL each) were used for measurement of TNF-α and IL-6 concentrations.

### Western blot

The ovarian tissues of the mutants and WT control were homogenized and prepared by RIPA lysis buffer (EMD Millipore, Billerica, MA). The supernatant was collected, and the concentration of proteins was determined by the BCA protein assay (Thermo Scientific). The extracted proteins (10 µg) were subjected to electrophoresis on 12% SDS-polyacrylamide gel and electro-transferred onto PVDF membranes. The membranes were blocked with 5% non-fat milk and incubated with the primary antibody overnight followed by washing with TBST and incubation with the secondary antibody for 2 h. The signals were detected by ECL Reagent (Thermo Scientific). The primary antibody used was anti-cleaved Caspase-3 (1:1000, #9661, Cell Signaling Technology, Beverly, MA) and the secondary antibody was HRP-linked anti-rabbit IgG antibody (1:500, #7074, Cell Signaling Technology).

### Fertility assay

The fecundity of female mutant fish was assessed by natural mating with the WT males. Four females of each genotype were used to mate with four WT males individually, and the ovulated eggs were counted within 3 h after spawning. The tests were repeated at 4-day interval and the averages of four fish were plotted for statistical analysis.

### Statistical analysis

All values were expressed as mean ± standard error of the mean (SEM). The data were analyzed by Student’s t-test, one-way ANOVA or Chi-square analysis with Prism 8 (GraphPad, San Diego, CA). Significance was set at P < 0.05.

## Results

### Establishment of mutant zebrafish lines for activin β subunits

Zebrafish has three activin β subunits, including two forms of βA (*inhbaa* and *inhbab*) and one βB (*inhbb*). To understand the functional importance of the activin system in zebrafish, we disrupted all three β subunit genes using the CRISPR/Cas9 method. We targeted *inhbaa* in exon 1, *inhbab* in exon 2, and *inhbb* in exon 1 within the open reading frame region and downstream of the translation start site to generate *inhbaa*, *inhbab*, and *inhbb* single mutants, respectively. For the *inhbaa* mutant, a 13-bp deletion was introduced in exon 1 (ZFIN line number: umo27) (Fig. S1Aa). To increase the efficiency of mutagenesis, two CRISPR/Cas9 sites in exon 2 of *inhbab* were selected, and we obtained a 4-bp deletion mutant (umo28) (Fig. S1Ba). For *inhbb*, a 7- bp insertion mutant line was established (umo29) (Fig. S1Ca). The indel mutations in all three mutants introduced a new premature stop codon to disrupt protein translation. We also verified the mutations at transcript level in the ovary by RT-PCR using mutation-specific primers (F2 and R1), which would amplify the control sequence (+) but not the mutant (-) or vice versa. For *inhbaa* and *inhbab*, the signals could be amplified in controls (+/+ and +/−) but not homozygous mutant (−/−) (Fig. S1Ab and Bb). For *inhbb*, the amplified signals could be detected in mutants (+/− and −/−), but not WT control (+/+) (Fig. S1Cb).

### Roles of activin subunits in zebrafish development

In addition to stimulating pituitary FSH biosynthesis, activin was also discovered to function as a potent mesoderm-inducing factor (MIF) in *Xenopus* development [5, 6]. However, this role has not been supported in the mouse model by genetic approach [8, 9]. To demonstrate whether activin is involved in mesoderm formation in fish, we examined the ratios of three genotypes (+/+, +/−, and −/−) in the offspring obtained from incrossing of the heterozygotes (+/− x +/−) for *inhbaa*, *inhbab* and *inhbb*. As shown in Fig. 1A, the distribution of three genotypes (+/+, +/−, and −/−) for *inhbab* and *inhbb* were normal according to the Mendelian Law (0.25:0.5:0.25) at 90 dpf or 3 months post-fertilization (mpf); however, the survival rate of *inhbaa* mutant was only about half of the expected level (13.6%), suggesting higher mortality in *inhbaa−/−* (Fig. 1A).

**Figure 1.**
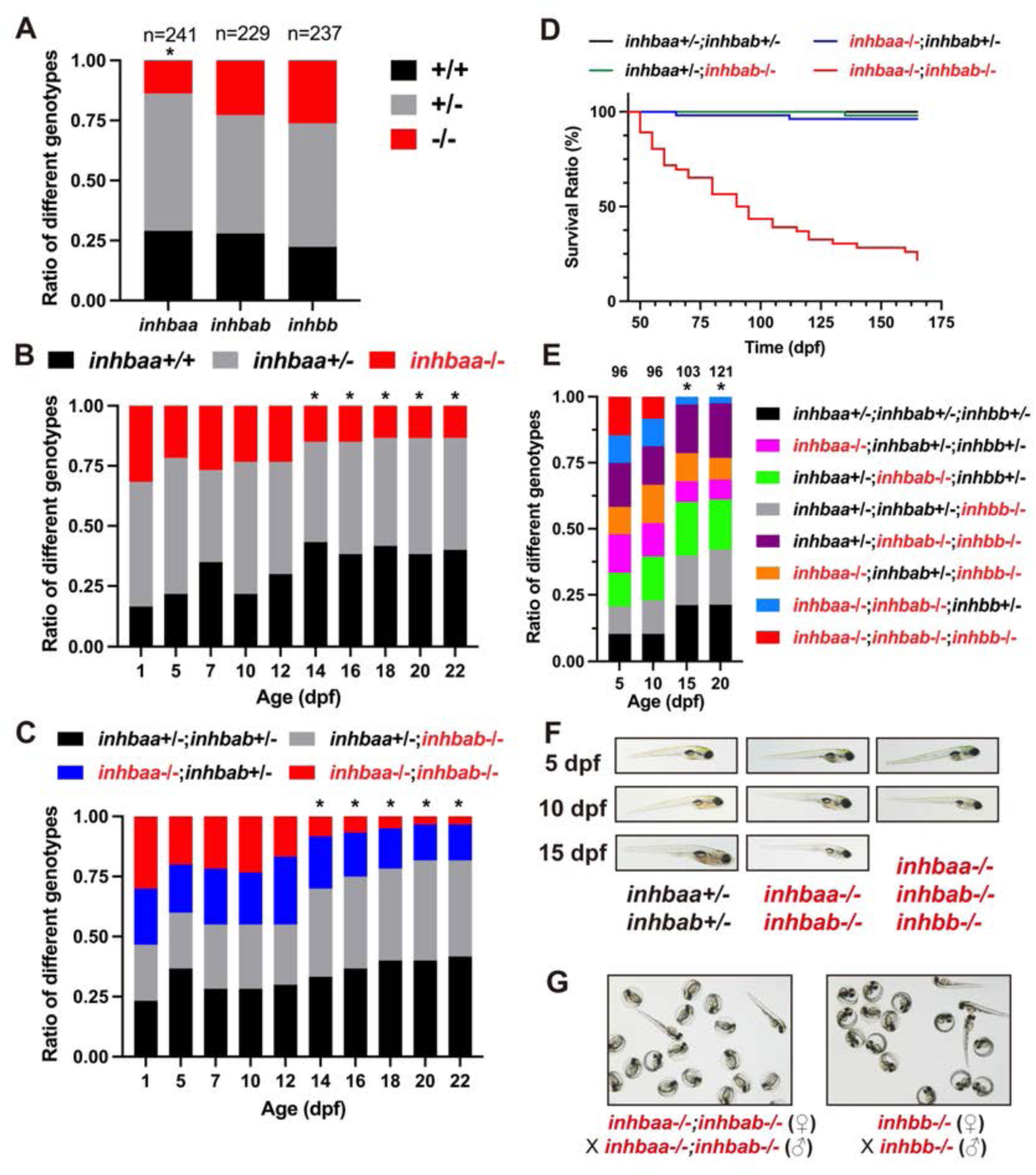
Analysis of survival ratios in activin β subunit mutants. (A) Ratios of different genotypes in the offspring of heterozygotes (+/− x +/−) at adult stage. (B) Ratios of different genotypes in the offspring of *inhbaa*+/− x *inhbaa*+/− from 1 to 22 dpf. Sixty fish were sampled at each time point (n = 60). (C) Ratios of different genotypes in the offspring of *inhbaa*+/−;*inhbab*−/− x *inhbaa*−/−;*inhbab*+/− from 1 to 22 dpf. Sixty fish were sampled at each time point (n = 60). (D) Kaplan–Meier plot for activin βA single and double mutants (50 to 165 dpf; n = 46-54). (E) Ratios of different genotypes in the offspring of *inhbaa*+/−;*inhbab*−/−;*inhbb*+/− (♀) x *inhbaa*−/−;*inhbab*+/−;*inhbb*−/− (♂) from 5 to 20 dpf. The number of fish examined is shown on top of each column. (F) The larvae of control (*inhbaa*+/−;*inhbab*+/−), βA double mutant (*inhbaa*−/−;*inhbab*−/−) and βA/B triple mutant (*inhbaa*−/−;*inhbab*−/−;*inhbb*−/−) from 5 to 15 dpf. No triple mutant individuals could survive to 15 dpf. (G) Offspring of *inhbaa*−/−;*inhbab*−/− (♀) x *inhbaa*−/−;*inhbab*−/− (♂) and *inhbb*−/− (♀) x *inhbb*−/− (♂) at 72 hpf. The asterisk indicates significant deviation from expected ratios by Chi-square analysis (p<0.05).

To explore when the mortality occurred in *inhbaa−/−* mutant, we analyzed temporal changes of the three genotypes (+/+, +/−, and −/−) at different time points from 1 to 22 dpf. The results showed normal ratios from 1 to 12 dpf. However, a sharp drop occurred in the homozygous mutant (*inhbaa*−/−) at 14 dpf and the ratio remained low but stable afterwards (Fig. 1B). To demonstrate if the two βA genes (*inhbaa* and *inhbab*) had compensatory effects, we generated a double homozygous mutant (*inhbaa*−/−;*inhbab*−/−) by crossing *inhbaa*+/−*;inhbab*−/− (♀) with *inhbaa*−/−*;inhbab*+/− (♂). A more severe larval death was found at 14 dpf in the double mutant compared with the single *inhbaa*−/− mutant, and its ratio declined progressively afterwards, reaching only 5% at 22 dpf (Fig. 1C). These double mutant fish continued to die towards adult stage with only a few individuals survived at 165 dpf, in contrast to the control (*inhbaa*+/−*;inhbab*+/−) and single mutants (*inhbaa*−/− and *inhbab*−/−) (Fig. 1D).

To provide further evidence for roles of activin in early development, we went on to generate a triple mutant of all three β genes (*inhbaa*−/−*;inhbab*−/−*;inhbb*−/−) by crossing *inhbaa*+/−*;inhbab*−/−*;inhbb*+/− (♀) with *inhbaa*−/−*;inhbab*+/−*;inhbb*−/− (♂). Interestingly, the ratio of the triple mutant was 14.6% at 5 dpf, close to the expected ratio of 12.5%; however, the ratio dropped to 8.3% at 10 dpf, and we could not detect any triple mutant fish at 15 and 20 dpf (Fig. 1E), suggesting post-larval lethality without activins. The larval fish were morphologically normal at 5 and 10 dpf for both βA double mutant (*inhbaa*−/−*;inhbab*−/−) and βA/B triple mutant (*inhbaa*−/−*;inhbab*−/−*;inhbb*−/−) (Fig. 1F), suggesting normal embryonic development, including the mesoderm, without activins. We also demonstrated that all single (*inhbaa*−/−*, inhbab*−/− and *inhbb*−/−) and double homozygous mutants (*inhbaa*−/−*;inhbab*−/−*, inhbaa*−/−*;inhbb*−/− and *inhbab*−/−*;inhbb*−/−) (♀−/− x ♂−/−) could produce normal offspring, ruling out the involvement of maternal activins (Fig. 1G). Unfortunately, we could not test the triple mutant due to its complete post-larval lethality.

### Localization of activin-inhibin-follistatin system in ovarian follicles

Having examined involvement of activins in embryonic and post-larval development, we then focused our analysis on roles of activins in reproduction, especially in females. We first examined the expression of activin subunits (*inhbaa*, *inhbab* and *inhbb*), activin antagonist inhibin (*inha*) and binding protein follistatins (*fsta* and *fstb*) by RT-PCR in the somatic follicle layer and oocyte at FG stage. We previously demonstrated that activin βAa (*inhbaa*), βB (*inhbb*) and inhibin α (*inha*) were exclusively expressed in the follicle layer while follistatin a (*fsta*) was expressed primarily in the oocyte [31, 55]. In this study, we expanded the analysis by including activin βAb (*inhbab*) and follistatin b (*fstb*) to generate a more complete picture about the spatial distribution of the activin-inhibin-follistatin system in the follicle. Consistent with our previous results, all activin subunits (*inhbaa, inhbab* and *inhbb*) were expressed exclusively in the somatic follicle layer together with *inha*. In contrast, *fsta* was exclusively expressed in the oocyte whereas *fstb* was expressed equally in both the oocyte and follicle cells (Fig. 2).

**Figure 2.**
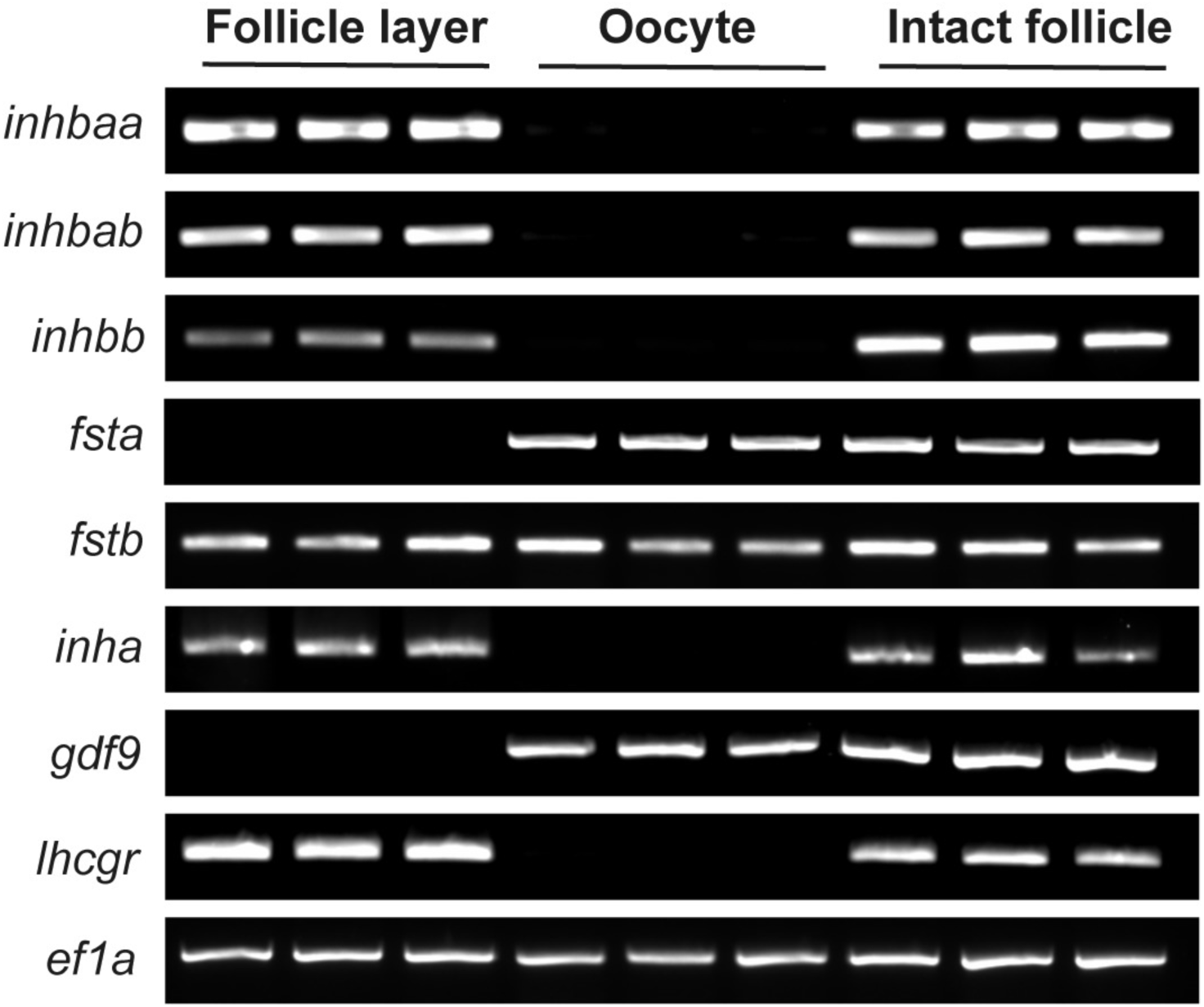
Intrafollicular distribution of activin-inhibin-follistatin system in full-grown (FG) follicles. The activin β subunits (*inhbaa*, *inhbab* and *inhbb*) and inhibin α (*inha*) were exclusively expressed in the follicle layer whereas follistatin a (*fsta*) was exclusively expressed in the oocyte. Follistatin b (*fstb*) was expressed in both follicle layer and oocyte. The housekeeping gene (*ef1a*) was expressed in both cell types, while *gdf9* and *lhcgr* were exclusively expressed in oocyte and follicle layer, respectively, indicating clean separation of the two compartments.

### Delayed follicle activation and puberty onset in activin βA-deficient females

Follicle activation is a critical event in fish folliculogenesis, which marks the transition from PG (stage I) to PV (stage II) stage, or primary growth to secondary growth (SG) phase. The SG phase begins with PV stage, which is characterized with the formation of cortical vesicles/alveoli in the oocyte. The first wave of PG-PV transition also marks puberty onset in female zebrafish, which occurs around 45 dpf when BL and BW reach 1.8 cm and 100 mg, respectively [56, 57]. The expression of *inhbaa* increased significantly during the PG-PV transition [32, 33], suggesting an important role for activin in the transition. To test this hypothesis, we examined females of activin βA mutants at 45 dpf. Unexpectedly, we did not see any obvious abnormalities in ovaries of the single mutants (*inhbaa*−/− and *inhbab*−/−) as compared to the control (*inhbaa*+/−*;inhbab*+/−). PV follicles appeared in the ovaries of both control and the single mutants once the BL and BW reached or exceeded the thresholds (1.8 cm/100 mg) for puberty onset. However, in contrast to the control and single mutants, the double mutant of the two activin βA subunits (*inhbaa*−/−*;inhbab*−/−) often showed a significant delay in puberty onset with absence of PV follicles in females whose body sizes were far beyond the thresholds (*eg*., 2.2 cm/138 mg) (Fig. 3).

**Figure 3.**
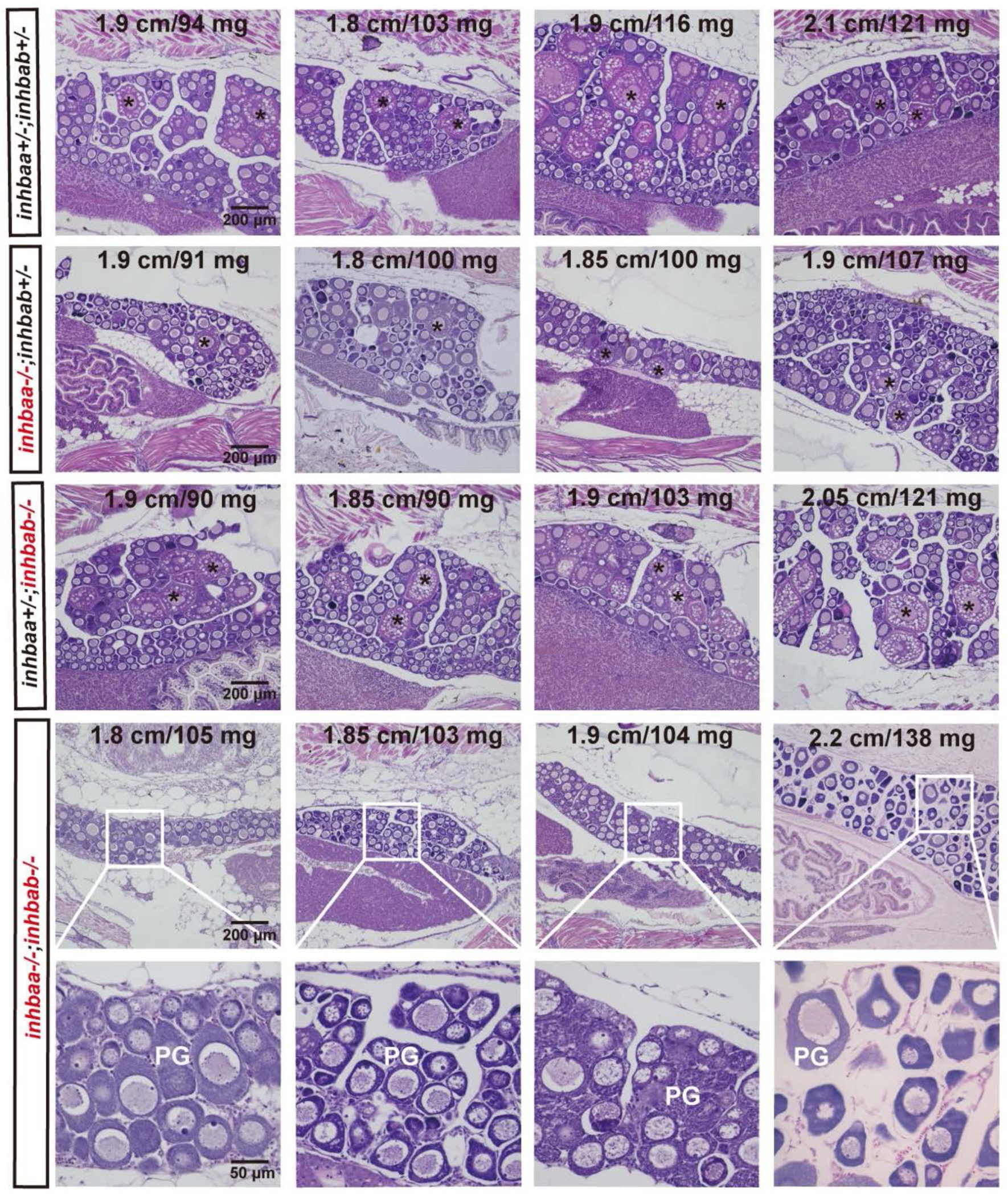
Delayed follicle activation and puberty onset in activin βA double mutant (*inhbaa*−/−;*inhbab*−/−) at 45 dpf. In control (*inhbaa+/−;inhbab+/−*) and single mutant (*inhbaa−/−*, *inhbab−/−*) fish, PV follicles with cortical alveoli in the oocyte started to appear when their BL and BW reached the threshold for puberty onset (1.80 cm/100 mg). However, in the double mutant (*inhbaa*−/−;*inhbab*−/−), PV follicles did not appear in many individuals although their body size had reached the threshold. The BL (cm) and BW (mg) of each fish are shown on the top. The asterisk shows PV follicles. PG, primary growth stage.

To further characterize the PG-PV transition, we divided the PV stage into three sub-stages: PV-I, PV-II and PV-III (Fig. 4A). PV-I is the starting stage when a single layer of small cortical alveoli appears in the ooplasm. In PV-II, the cortical alveoli remain as a single layer, but they become much larger. Multiple layers of large cortical alveoli are present in PV-III follicles. Many double mutant fish had PG follicles only even though their body size had exceeded the thresholds (1.8 cm/100 mg). For those with PV follicles, the follicles were mostly at PV-I stage, exhibiting an obvious delay in PG-PV transition and subsequent follicle growth (Fig. 4B). Quantitative analysis showed that the delayed follicle activation might also occur in *inhbaa*−/− single mutant, but it was not significantly different from the control and *inhbab*−/−. The delay was significant in the double mutant (*inhbaa*−/−*;inhbab*−/−) compared to the control and two single mutants (Fig. 4C). Consistently, analysis of follicle composition at 45 dpf showed a significantly decreased ratio of PV follicles in the double mutant compared to the control and two single mutants, supporting the view that there was a delayed exit of follicles from PG pool to PV stage in activin βA-deficient females (Fig. 4D).

**Figure 4.**
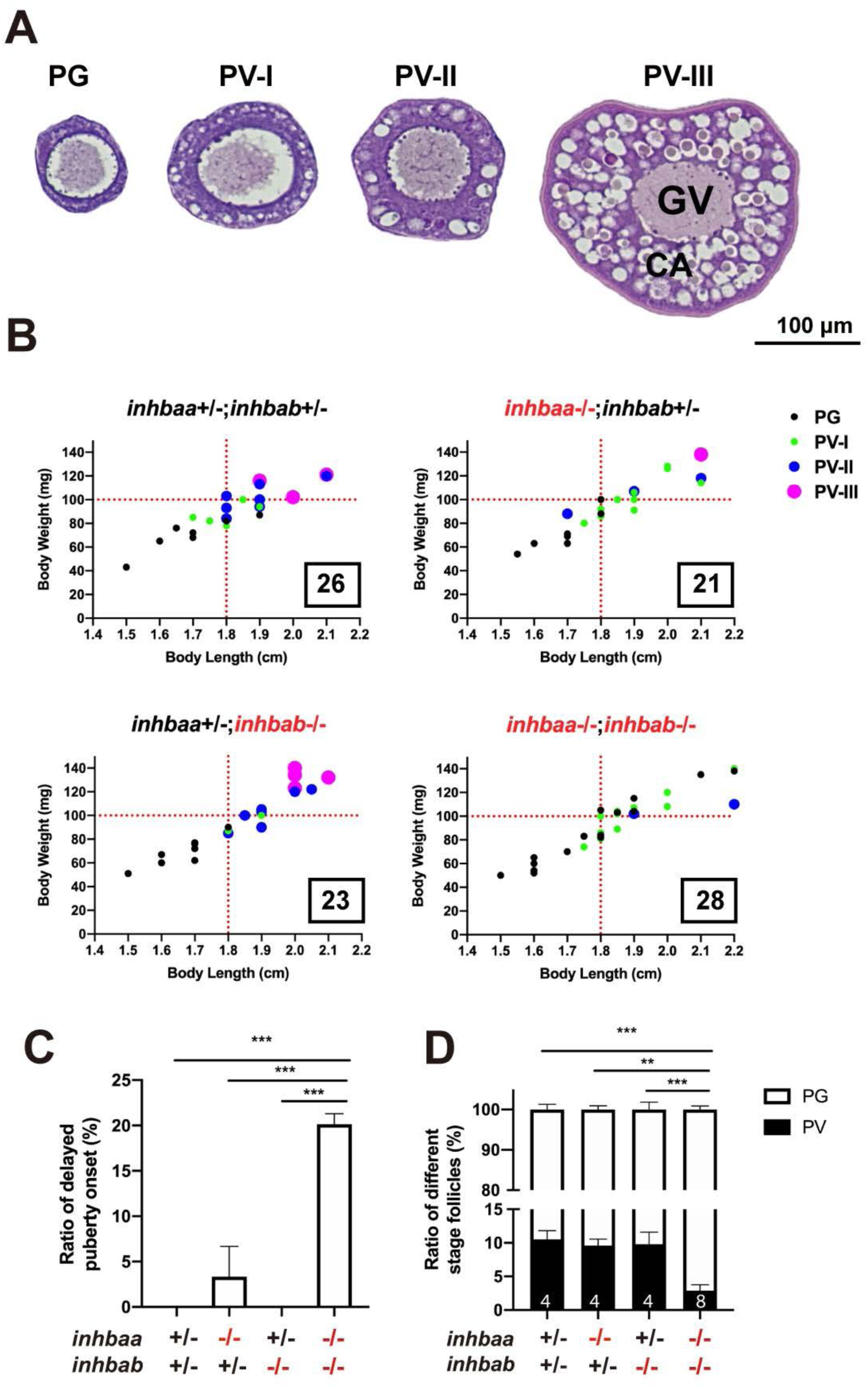
Quantitative analysis for delayed puberty onset in activin βA double mutant (*inhbaa*−/−;*inhbab*−/−). (A) Classification of PV follicles. The PV stage is further divided into three sub-stages according to size and layer number of the cortical alveoli (CA). PV-I, single layer of small CA; PV-II, single layer of large CA; PV-III, multiple layers of large CA. (B) Correlation between PG-PV transition and body size [BL (cm) and BW (mg)]. Dots in different color and size represent different stages of follicles. The number in the box indicates the sample size. (C) Ratios of delayed puberty onset in different activin βA mutants at 45 dpf (n = 3 batches). (D) Follicle composition in the ovary of βA mutants at 45 dpf. Sample size is indicated at the bottom of each column. (**P < 0.01; ***P < 0.001). PG, primary growth; PV, previtellogenic; GV, germinal vesicle.

### Defective post-pubertal follicle development in βA-deficient females

To investigate if activin deficiency had any effect on post-pubertal folliculogenesis, we examined young females at different time points of sexual maturation from puberty onset to maturity (55, 60, 70 and 80 dpf). The control and two single βA mutants (*inhbaa*−/− and *inhbab*−/−) exhibited normal follicle development with different stages of follicles present in the ovary, including PG, PV, EV, MV, LV and FG. Despite the delay in follicle activation or PG-PV transition, the subsequent vitellogenic growth could proceed normally in most individuals of βA double mutant (*inhbaa*−/−*;inhbab*−/−). However, the double mutant ovaries started to show signs of follicle degeneration or atresia after 60 dpf with increased inter-follicular spaces. In addition, we frequently observed hypertrophic granulosa cell layer in the double mutant follicles, which partially phenocopied the mutant of inhibin (*inha*) [45] (Fig. 5). Interestingly, some βA double mutant females (∼10%) could never start vitellogenic growth (70-135 dpf), and their ovaries contained mostly degenerating PG and sometimes a few PV follicles, which were separated by abundant somatic cells infiltrating the inter-follicular spaces (Fig. 6).

**Figure 5.**
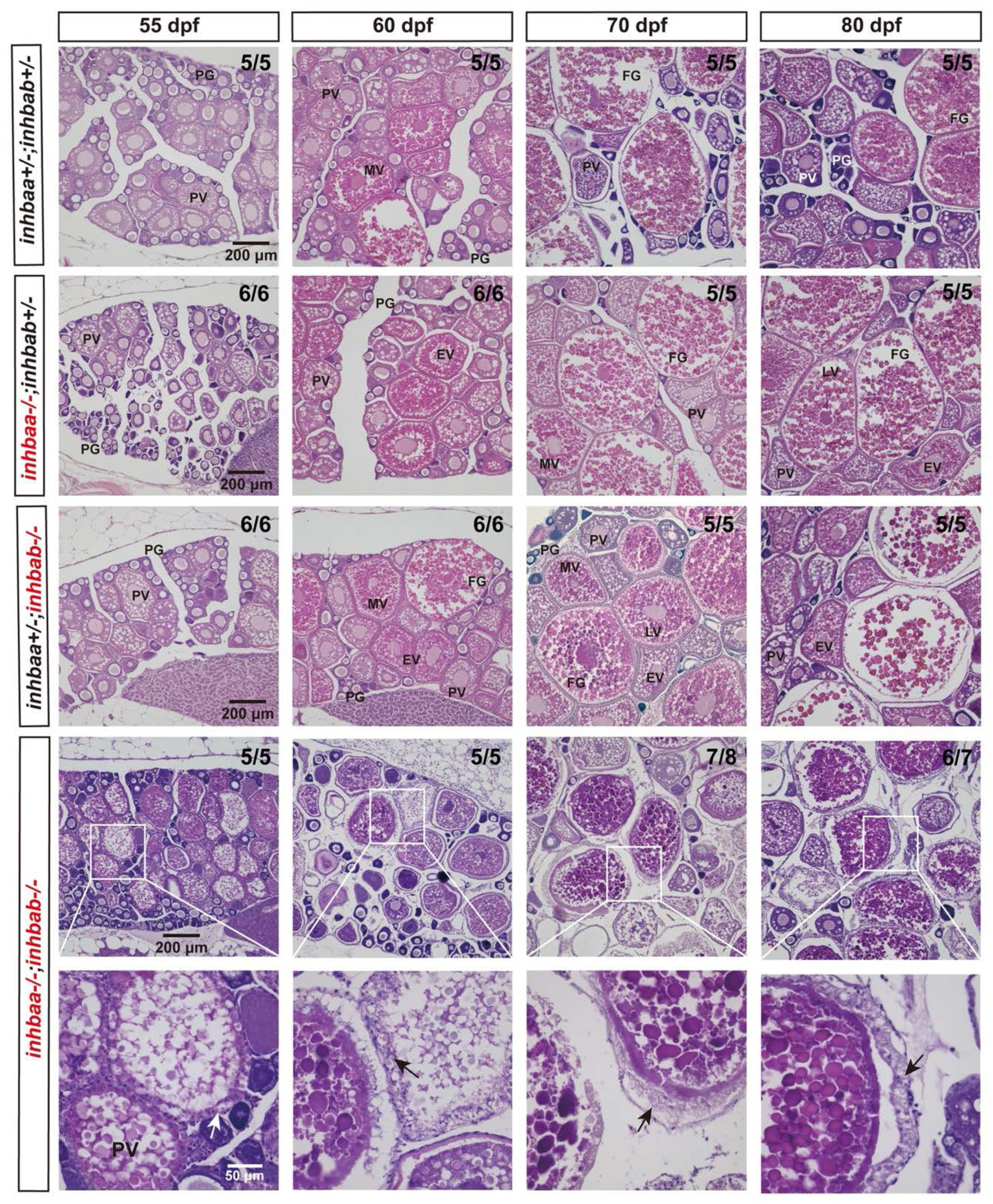
Histological analysis of activin βA mutant females during post-pubertal maturation (55-80 dpf). Follicles from control and βA single mutants (*inhbaa−/−*, *inhbab−/−*) showed normal growth and development. In the double mutant (*inhbaa*−/−;*inhbab*−/−), abnormal follicles with hypertrophic granulosa cells could often be observed (arrow). The boxed areas are shown at higher magnification below. The numbers shown in the photos indicate the total number of fish examined (lower) and the fish exhibiting similar phenotype to that shown (upper). PG, primary growth; PV, previtellogenic; EV, early vitellogenic; MV, mid-vitellogenic; LV, late vitellogenic; FG, full-grown.

**Figure 6.**
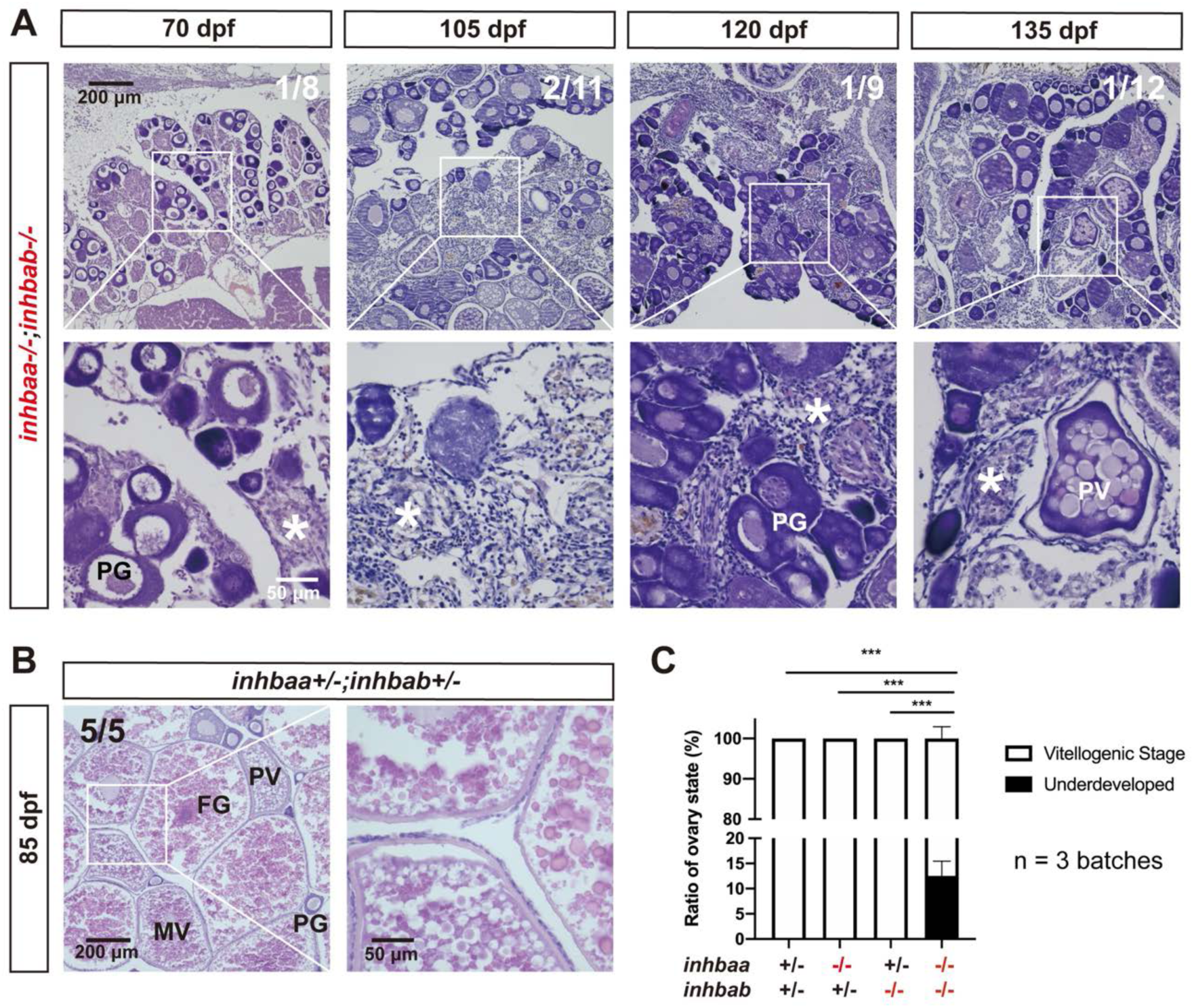
Failure of follicle activation and defective folliculogenesis in activin βA double mutant (*inhbaa*−/−;*inhbab*−/−). (A) Ovary of some mutant individuals (∼10%) at 70-135 dpf. The ovary contained PG follicles only at 70 dpf, and much of the inter- follicular space was filled with abundant stromal cells. At later development stages (105, 120 and 135 dpf), PV follicles could occasionally be observed; however, there were no vitellogenic follicles and the ovaries were mostly occupied by over-proliferated stromal cells. The boxed areas are shown at higher magnification below. The numbers shown in the photos indicate the total number of fish examined (lower) and the fish exhibiting similar phenotype to that shown (upper). (B) Ovary in the control fish (*inhbaa*+/−*;inhbab*+/−) at 85 dpf. Vitellogenic follicles developed normally in the control. (C) Statistical analysis of ovarian types from 105 to 135 dpf (***p < 0.001; n = 3 batches). PG, primary growth; PV, previtellogenic; MV, mid-vitellogenic; FG, full-grown; asterisk, infiltrating stromal tissue.

### Defective ovarian maintenance in βA/B-deficient females

Despite the defective phenotypes observed in activin βA double mutant females (*inhbaa*−/−*;inhbab*−/−) in terms of puberty onset and post-pubertal follicle development, the single mutants of all three β subunits (*inhbaa*, *inhbab* and *inhbb*) were largely normal up to sexual maturity and the follicles in most βA double mutants could also proceed beyond PV stage to FG stage. To investigate long-term functions of activin subunits in folliculogenesis, we examined all three single mutants (*inhbaa−/−*, *inhbab−/−* and *inhbb−/−*) and βA double mutant (*inhbaa*−/−*;inhbab*−/−) at different time points after maturation (90, 120, 180, 240, 270 and 300 dpf).

At 90 dpf, the control (*inhbaa*+/−*;inhbab*+/−) and two βA single mutant females (*inhbaa*−/− and *inhbab*−/−) were normal at histological level with all stages of follicles present in the ovary without any abnormalities. However, in the ovary of βA double mutant (*inhbaa*−/−*;inhbab*−/−), the stromal cells were abundant, filling up the spaces between follicles (Fig. 7A). The double mutant also seemed to grow more slowly as its BL, but not BW, was significantly lower than the control and single mutants with high variation (Fig. 7B). Quantification of follicle composition at 90 dpf showed that the double mutant had significantly more PG follicles but less PV and vitellogenic follicles (EV-FG) compared to the control and single mutants, in agreement with the observation that the loss of βA subunits caused a delay in follicle activation or PG-PV transition (Fig. 7C).

**Figure 7.**
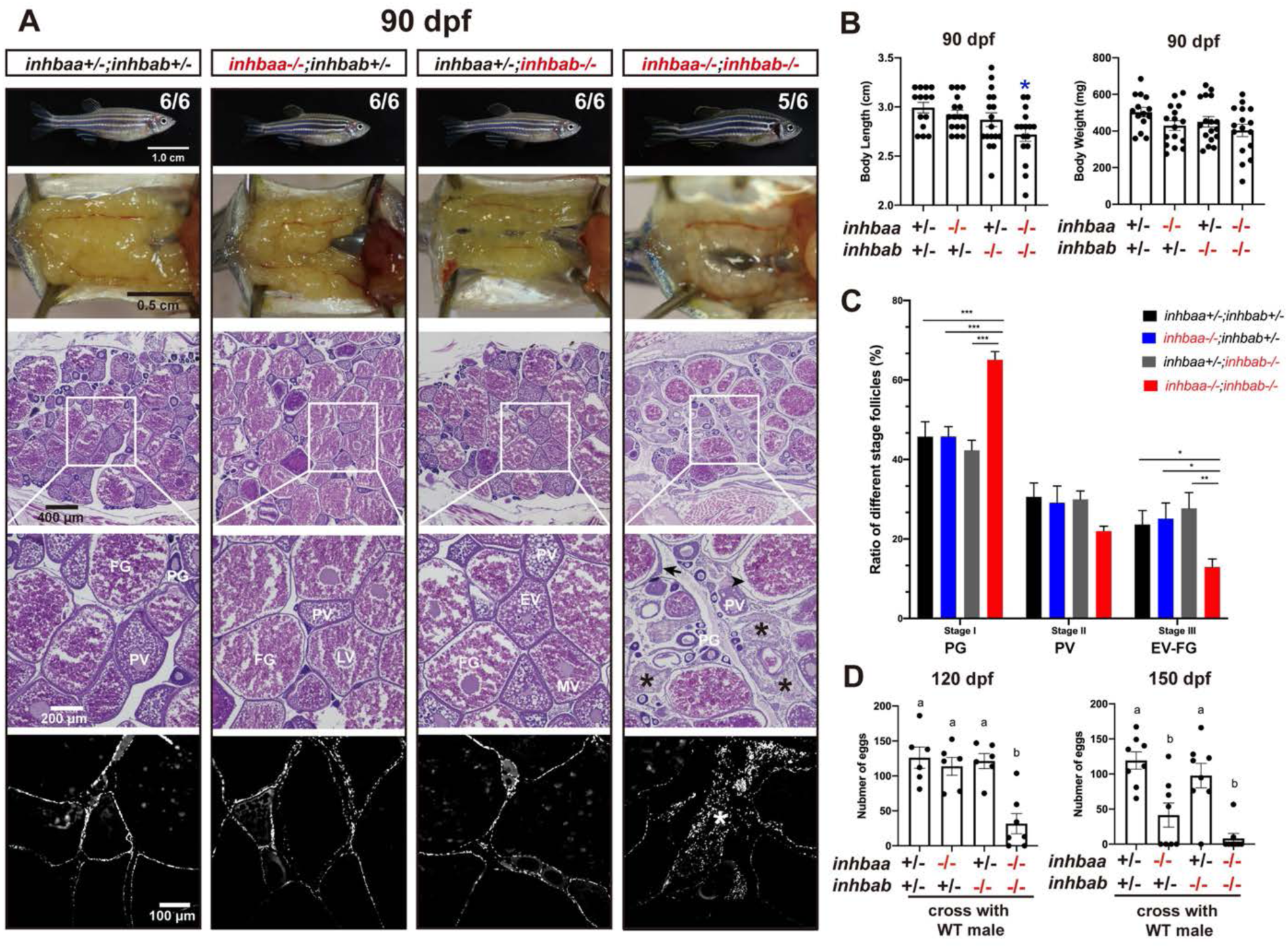
Follicle development and fecundity in young and mature females of βA mutants. (A) Morphology, gross anatomy and histological analysis of activin βA female mutants at 90 dpf. Follicles from the control (*inhbaa*+/−*;inhbab*+/−) and single mutants (*inhbaa−/−*, *inhbab−/−*) showed normal growth and ovarian development. The double mutant (*inhbaa*−/−;*inhbab*−/−) started to show stromal cell hyperplasia (asterisk), granulosa cell hypertrophy (arrow) and follicle atresia (arrowhead). The overproliferation of the stromal cells is also shown by DAPI staining for nuclei (bottom). The boxed areas are shown at higher magnification below. The numbers shown in the photos indicate the total number of fish examined (lower) and the fish exhibiting similar phenotype to that shown (upper). (B) Body length and body weight of activin βA mutants at 90 dpf. (C) Follicle composition in the ovaries of the activin βA mutants at 90 dpf (n = 5). (*p < 0.05; **p < 0.01; ***p < 0.001). PG, primary growth (stage I); PV, previtellogenic (stage II); EV-FG, vitellogenic (stage III). (D) Fertility and fecundity of activin βA female mutants at 120 and 150 dpf. The mutant females were bred with normal WT males by natural breeding. Different letters indicate statistical significance (p < 0.05). Each dot represents the average egg number of four females from each spawning test (n = 6 tests).

At 120 dpf, the βA double mutant females (*inhbaa*−/−;*inhbab*−/−) showed a significantly reduced fecundity compared with the control and single mutants. Interestingly, the fecundity of *inhbaa* single mutant (*inhbaa*−/−) also started to decrease significantly at 150 dpf when the double mutant became almost infertile (Fig. 7D). Histological analysis at 120 dpf demonstrated stromal cell hyperplasia and less vitellogenic follicles in the βA double mutant. Although most follicles of *inhbaa* single mutant appeared normal, we often observed degenerative vitellogenic follicles near the surface of the ovarian lamellae. By comparison, the ovaries of control and *inhbab* single mutant were normal (Fig. 8A). At 180 dpf, both *inhbaa* single mutant and double βA mutant had lost fertility and their ovaries became more degenerative with increasing somatic cells and atretic follicles (Fig. 8B). Due to low survival rate of the βA double mutant, we only had chance to examine the single mutants of *inhbaa* and *inhbab* at later stages. At 240 and 300 dpf, the somatic cells continued to proliferate and they occupied vast areas in the *inhbaa−/−* ovary whereas the ovary and folliculogenesis in *inhbab−/−* single mutant remained normal (Fig. 9). Examination of *inhbaa−/−* females at 18 mpf (> 540 dpf) showed severe defects in ovarian structure. The mutant ovaries had all regressed, becoming barely visible to eyes in some individuals. Histological examination demonstrated that folliculogenesis had ceased in all mutant fish examined (8 in total) with follicles arrested mostly or completely at PG stage and most parts of the ovaries were occupied by stromal cells or amorphic fibrous tissues. Interestingly, all *inhbaa−/−* fish examined at this time point developed spine curvature to different degrees (Fig. 10).

**Figure 8.**
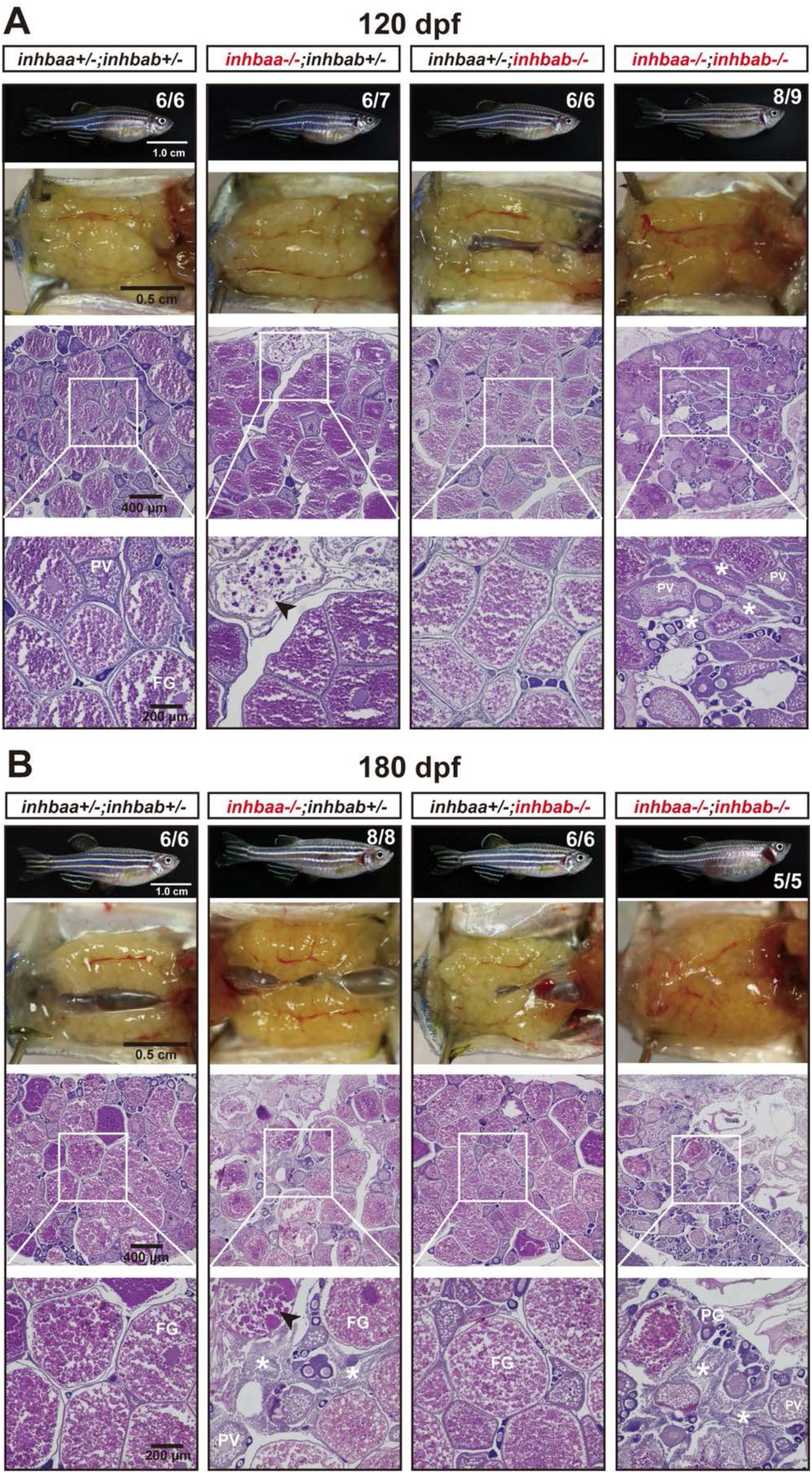
Ovaries of activin βA female mutants at 120 dpf (A) and 180 dpf (B). Follicles in the control (*inhbaa*+/−;*inhbab*+/−) and *inhbab* single mutant (*inhbab*−/−) were normal. The *inhbaa* single mutant (*inhbaa*−/−) and double mutant (*inhbaa*−/−;*inhbab*−/−) often contained degenerative or atretic vitellogenic follicles (arrowhead) and abundant stromal cells in the inter-follicular space (asterisk). The boxed areas are shown at higher magnification below. The numbers shown in the photos indicate the total number of fish examined (lower) and the fish exhibiting similar phenotype to that shown (upper).

**Figure 9.**
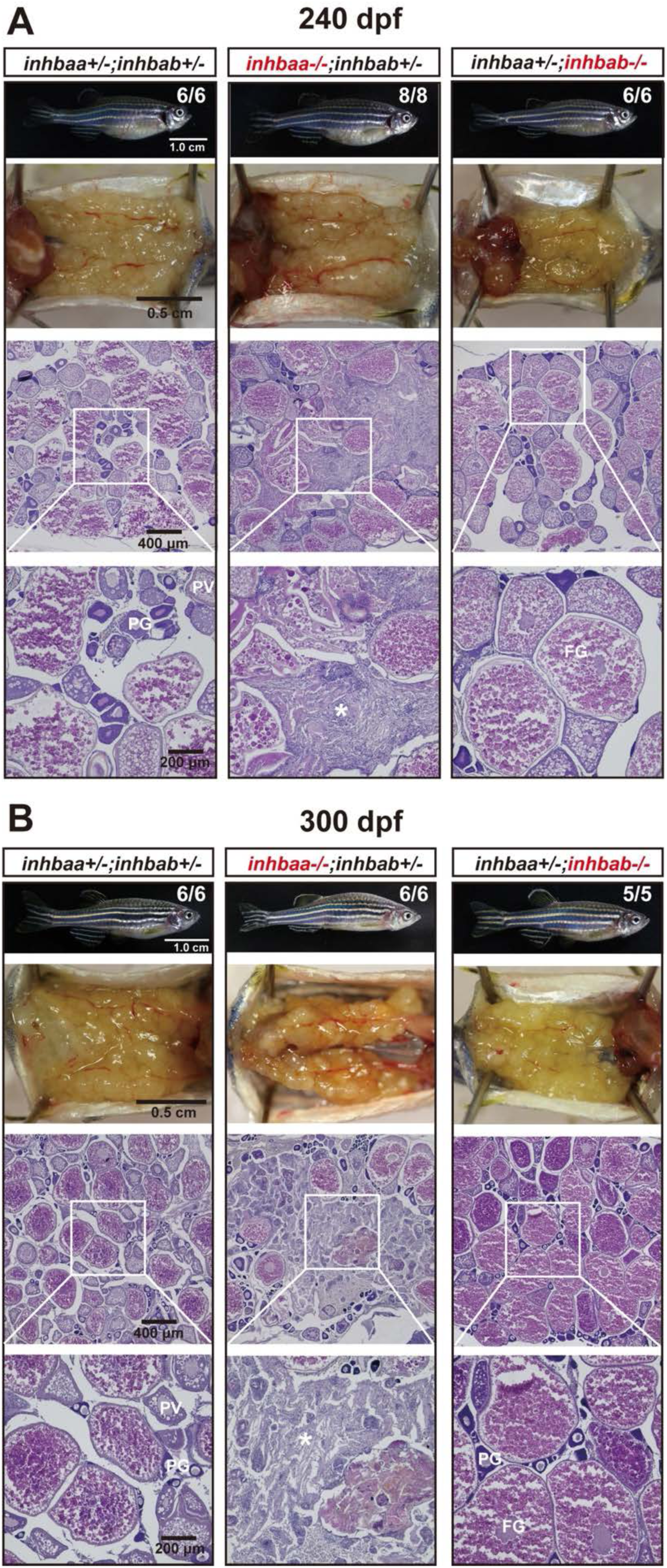
Ovaries of activin βA female mutants at 240 dpf (A) and 300 (B) dpf. The control and *inhbab* single mutant fish showed normal ovarian morphology and structure whereas the *inhbaa* mutant showed severe ovarian disorganization and dysfunction. Much of the space in the *inhbaa−/−* ovary was occupied by stromal cells and fibrous tissues (asterisk). The boxed areas are shown at higher magnification below. The numbers shown in the photos indicate the total number of fish examined (lower) and the fish exhibiting similar phenotype to that shown (upper). PG, primary growth; PV, pre-vitellogenic; FG, full-grown.

**Figure 10.**
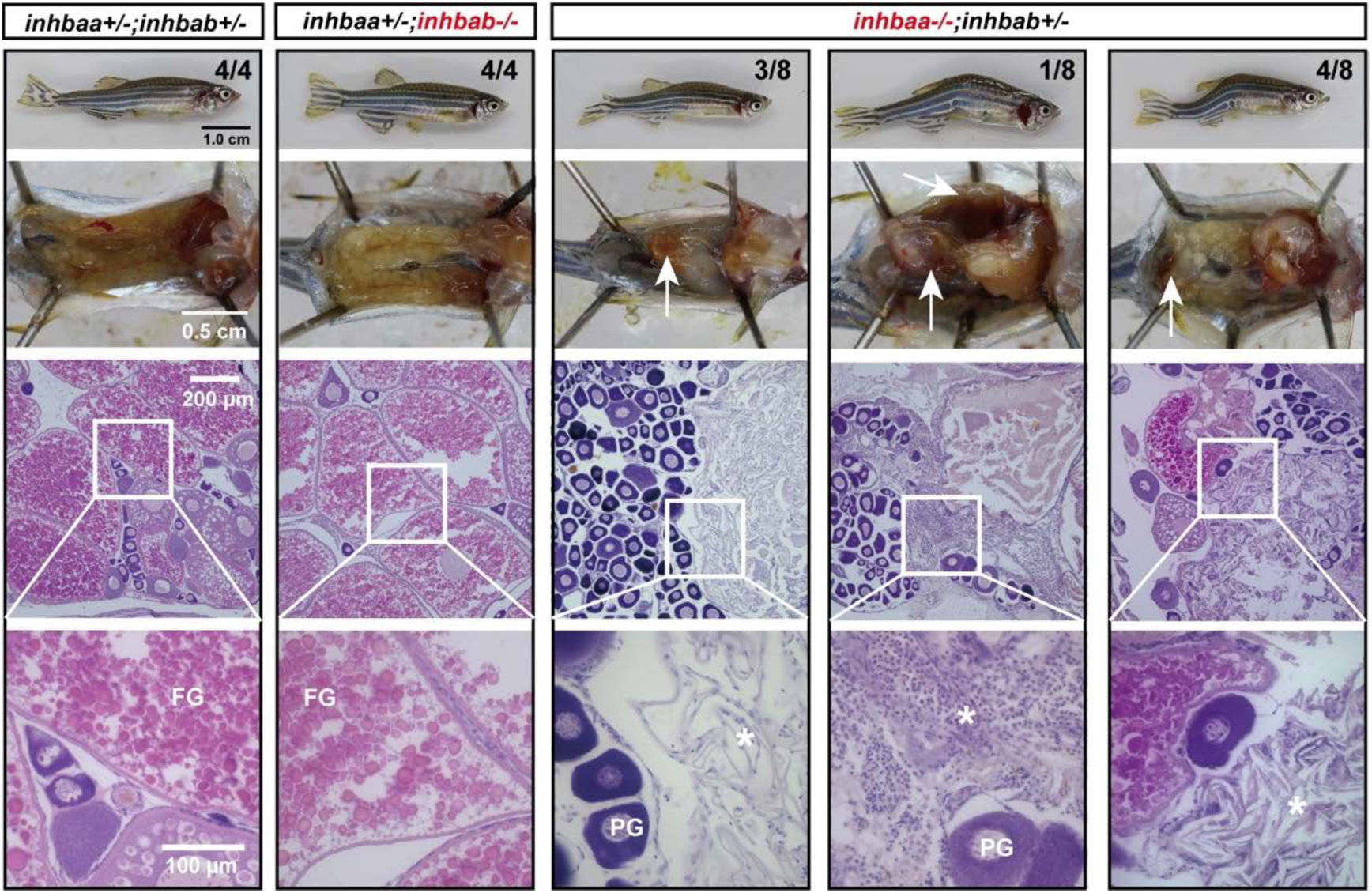
Long-term effect of *inhbaa* deficiency on ovarian maintenance. Tumour/cyst-like tissues (arrow) accumulated in the ovary of all individuals examined at ∼18 mpf (> 540 dpf). The folliculogenesis had ceased with follicles arrested at PG stage and the ovaries contained abundant stromal cells and fibrous tissues (asterisk). The control and *inhbab−/−* fish showed normal ovarian structure. The boxed areas are shown at higher magnification below. The numbers shown in the photos indicate the total number of fish examined (lower) and the fish exhibiting similar phenotype to that shown (upper). PG, primary growth; FG, full-grown.

In contrast to βA deficiency, the loss of βB subunit (*inhbb*−/−) did not generate any abnormal phenotypes up to maturity. The folliculogenesis was normal with no significant difference from the control fish from 45 to 90 dpf (Fig. 11A). At 180 dpf when βA-defficient females had lost fertility with severe ovarian dysfunction, the βB mutant females (*inhbb−/−*) were largely normal in both ovarian structure and fertility (fecundity) (Fig. 11B and C). However, the fecundity of the *inhbb−/−* mutant females decreased sharply at 270 dpf (Fig. 11C) and their ovaries contained significantly more PV follicles but less vitellogenic follicles (EV-FG) albeit without statistical significance (Fig. 11B and D), suggesting a possible blockade at PV-EV transition. Alternatively, the higher number of PV follicles might also be due to increased degeneration of the vitellogenic follicles in the mutant. In addition, there was an accumulation of extracellular fluid in the inter-follicular spaces, which were also infiltrated by abundant stromal cells (Fig. 11B).

**Figure 11.**
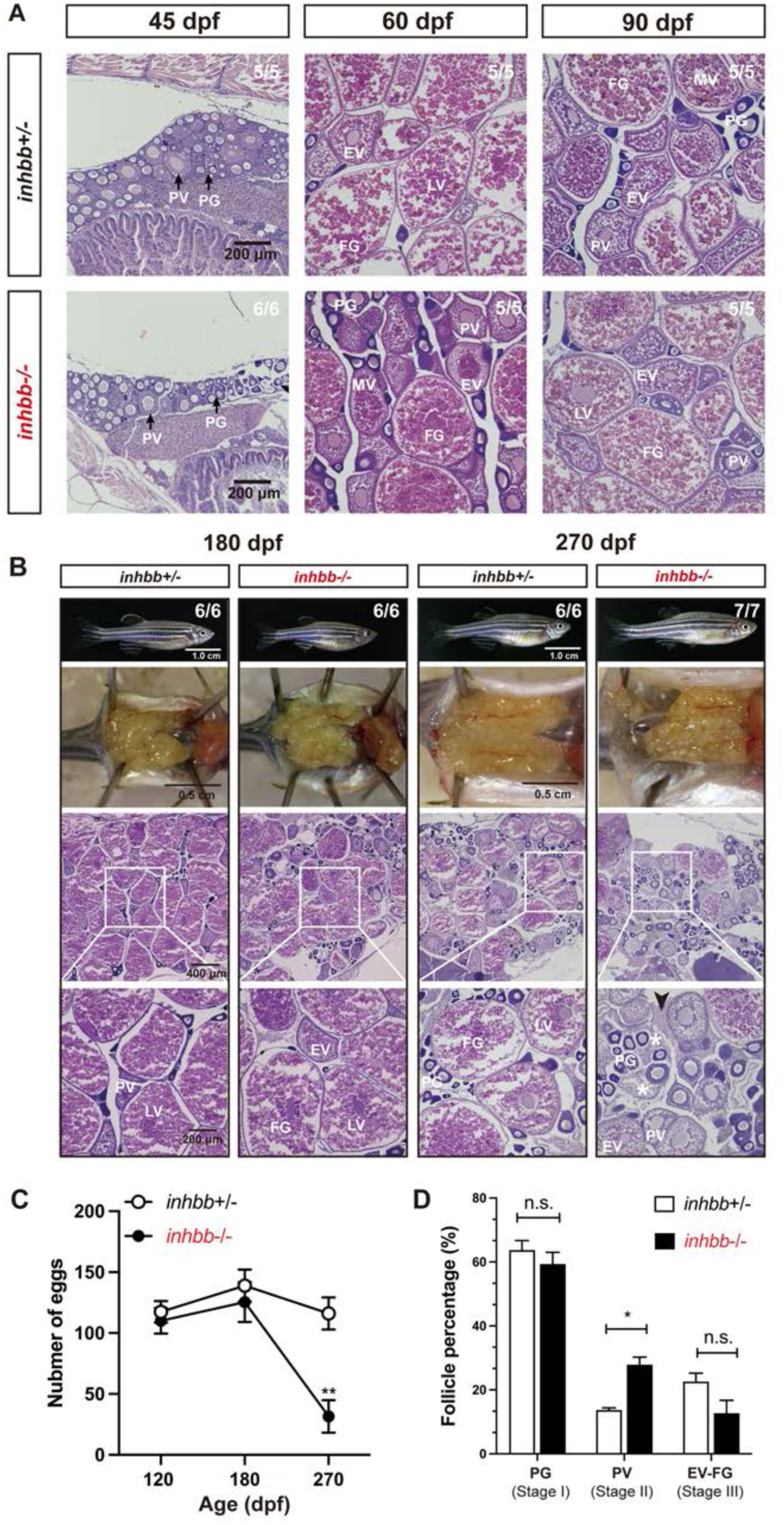
Phenotype analysis of activin βB mutant females at different stages (45-270 dpf). (A) Histology of ovaries from 45 to 90 dpf. The βB mutant (*inhbb*−/−) showed no abnormalities in follicle growth and composition compared with the control (*inhbb*+/−). (B) Histology of ovaries at 180 and 270 dpf. The mutant ovary remained normal at 180 dpf; however, the condition started to deteriorate at 270 dpf with significant changes in follicle composition and accumulation of stromal cells (asterisk) and fluid (arrowhead) between follicles. The numbers shown in the photos indicate the total number of fish examined (lower) and the fish exhibiting similar phenotype to that shown (upper). (C) Fecundity of control (*inhbb*+/−) and mutant (*inhbb*−/−) females at different times (120, 180 and 270 dpf). The female fish were bred with normal WT males by natural breeding, and the number of eggs released by each fish was counted and analyzed (**p < 0.01, n = 4). The fecundity of mutant females (*inhbb*−/−) dropped sharply at 270 dpf. (D) Follicle composition in the control (*inhbb*+/−) and mutant (*inhbb*−/−) ovaries at 270 dpf. The mutant fish contained more PV follicles but less vitellogenic follicles (EV-FG) at 270 dpf (*P < 0.05, n = 4 for *inhbb*+/− and 5 for *inhbb*−/−). PG, primary growth; PV, pre-vitellogenic; EV, early vitellogenic; MV, mid-vitellogenic; LV, late vitellogenic; FG, full-grown; n.s., no significance.

We also created double mutants of *inhbb*−/− together with *inhbaa−/−* or *inhbab−/−* (*inhbaa−/−;inhbb*−/−, *inhbab−/−;inhbb*−/−). For double mutant of *inhbaa* and *inhbb* (*inhbaa−/−;inhbb*−/−), we performed histological examination at 100, 165 and 240 dpf (Fig. 12A) and analyzed follicle composition at 165 dpf (Fig. 12B). The *inhbaa* single mutant (*inhbaa−/−;inhbb+/−*) and double mutant (*inhbaa−/−;inhbb−/−*) showed similar follicle composition with significantly more PG follicles but less vitellogenic follicles (EV-FG) compared to control (*inhbaa+/−;inhbb*+/−) and *inhbb* single mutant (*inhbaa+/−;inhbb*−/−) (Fig. 12B). As for the double mutant *inhbab−/−;inhbb−/−,* we examined the mutants at 100, 180 and 270 dpf (Fig. 12C) and analyzed follicle composition at 270 dpf (Fig. 12D). The double mutant *inhbab−/−;inhbb−/−* had similar follicle composition to that of the *inhbb−/−* single mutant, *i.e.,* more PV but less vitellogenic follicles (EV-FG). These observations indicate that the two double mutants phenocopied the single mutants of *inhbaa−/−* and *inhbb*−/−, respectively, in both timing and severity of the abnormalities, suggesting no functional compensation between βA and βB subunits.

**Figure 12.**
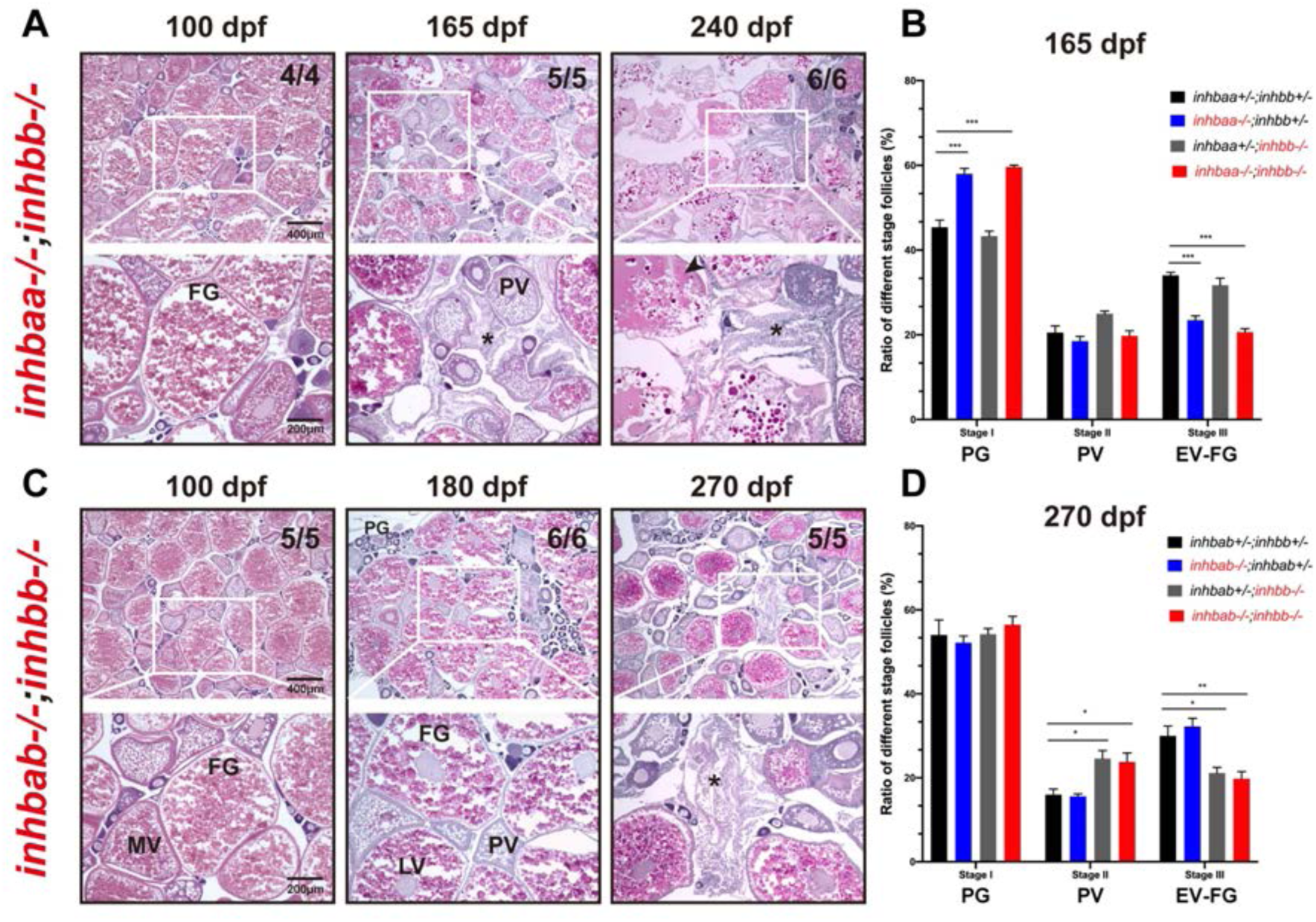
Phenotype analysis of βA and βB double mutants. Double mutants involving both βA and βB subunits were created (*inhbaa−/−;inhbb−/−*, and *inhbab−/−;inhbb−/−*) and their ovaries examined at different time points (100-270 dpf). (A) Histological analysis of *inhbaa−/−;inhbb−/−* from 100 to 240 dpf. (B) Follicle composition in the ovaries of the *inhbaa;inhbb* mutants at 165 dpf (*** p < 0.001, n = 4). (C) Histological analysis of *inhbab−/−;inhbb−/−* from 100 to 270 dpf. (D) Follicle composition in the ovaries of the *inhbab;inhbb* mutants at 270 dpf (* p < 0.05; ** p < 0.01, n = 4). No interactive effects were observed between βA and βB in terms of histological structure and follicle composition. The double mutants phenocopied the single mutants *inhbaa−/−* and *inhbb−/−*, respectively, showing similar ovarian disorganization, follicle degeneration, and accumulation of somatic stromal cells (asterisk) and fluid (arrowhead) at 165 and 240 dpf for *inhbaa−/−;inhbb−/−* and 270 dpf for *inhbab−/−;inhbb−/−*. The numbers shown in the photos indicate the total number of fish examined (lower) and the fish exhibiting similar phenotype to that shown (upper). PV, pre-vitellogenic; MV, mid-vitellogenic; LV, late vitellogenic; FG, full-grown.

### Potential role of activin βA in ovarian tumor/cyst formation, fibrosis and inflammation

In mammals and teleosts, inhibin α (INHA/Inha) has been reported to function as a tumor suppressor [45, 58]. We also investigated potential tumorigenic roles of activin β subunits in zebrafish. Similar to zebrafish *inha* mutant [45], tumors or cysts formed randomly in the ovary of about 40% *inhbaa−/−* mutant females after one year post-fertilization (> 12 mpf). The mutant ovary often displayed clumpy surface with numerous cysts or tumor-like outgrowths. Dissection of the ovaries often revealed brownish tissues of different sizes. Histological sectioning demonstrated degenerating follicles, abundant stromal cells and fiber-rich connective tissues (fibrosis) (Fig. 13A). TUNEL staining suggested active apoptotic activity among the stromal cells in the ovary of *inhbaa−/−* mutant, but not *inhbab−/−* (Fig. 13B) and this was confirmed by Western blot analysis showing increased level of cleaved Caspase-3 in *inhbaa−/−* ovary as compared to the control and *inhbab−/−* (Fig. 13C). Sirius red (SR) and Masson trichrome (MT) staining showed presence of abundant collagen fibers in the *inhbaa−/−* ovary, indicating ovarian fibrosis (Fig. 13D). In addition, measurement with ELISA showed increased levels of two proinflammatory cytokines, TNF-α and IL-6, in the ovary of *inhbaa−/−* mutant compared to the control and *inhbab−/−* mutant (Fig. 13E). We also determined expression levels of some functional genes implicated in apoptosis (*casp3a* and *p53*), ovarian tumors (*pawr*, *pax8*, *pgr*, *erbb2* and *cdkn2a/b*) [59–64] and fibrosis (*tgfb1a* and *pparg*) [65–68], and the results showed increased expression for most of these genes in *inhbaa−/−* ovary with the increase of *casp3a*, *pax8* and *tgfb1a* expression being statistically significant compared to the control and *inhbab−/−* mutant (Fig. 13F).

**Figure 13.**
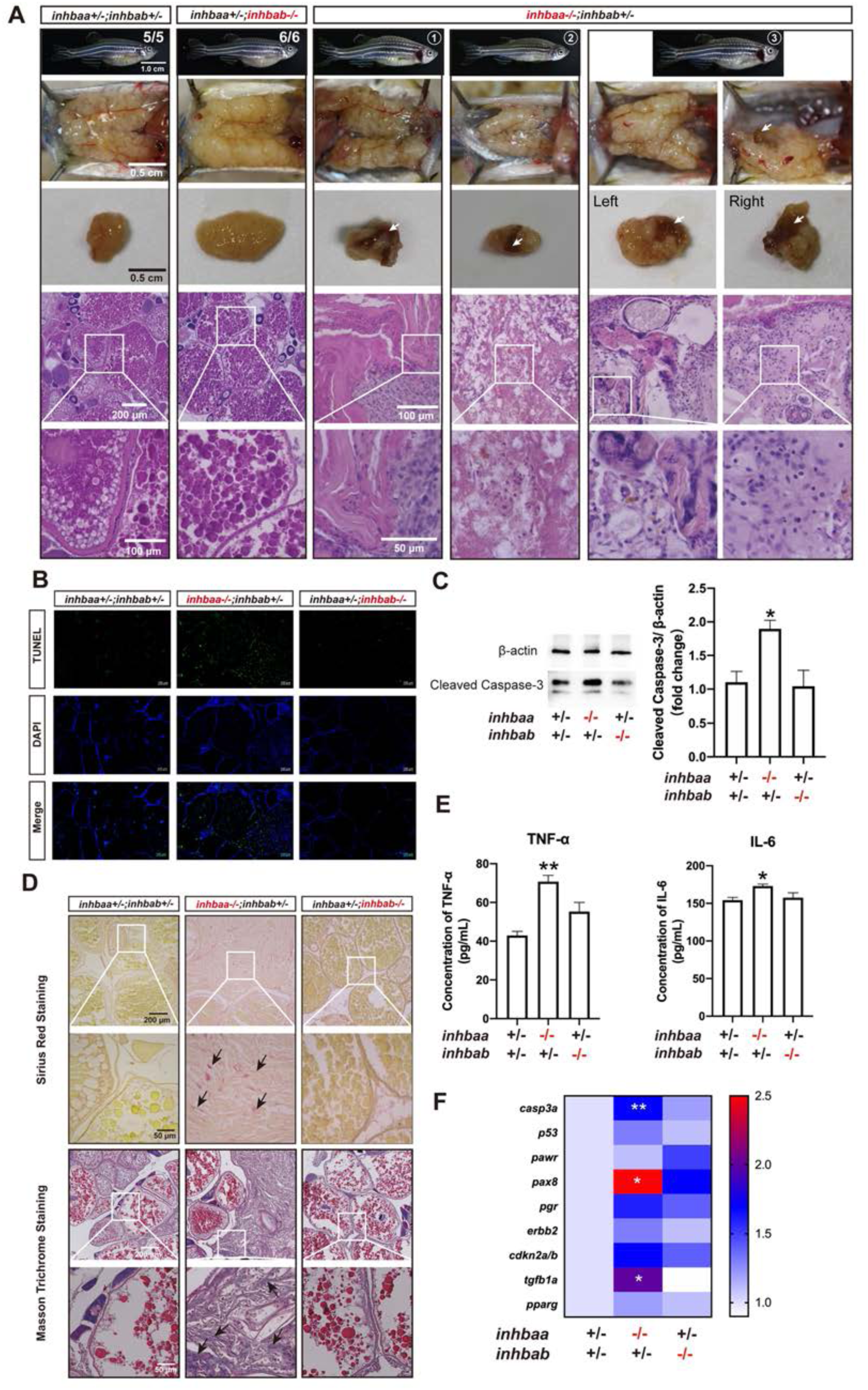
Ovarian disorders in aged *inhbaa*−/− mutant. (A) Tumorigenesis in mutant ovaries (>12 mpf). Tumor-like tissues or cysts accumulated in the mutant ovary unilaterally or bilaterally around one-year post-fertilization in all individuals examined (7 in total), and 3 fish contained brownish tissues of different sizes (arrow) (3/7). The fish on the right had the brown tissues on both ovaries. The boxed areas are shown at higher magnification below. (B) TUNEL staining for apoptosis in the ovary. Large areas of somatic cells in the ovary of *inhbaa−/−* but not *inhbab−/−* mutant showed strong TUNEL signals at 240 dpf. (C) Western blotting analysis for cleaved Caspase-3 in the ovary of βA single mutants (*inhbaa−/−*, *inhbab−/−*) at 240 dpf (n = 3). (D) Sirius red and Masson trichrome staining of the ovarian tissues in different genotypes of the βA mutants at 240 dpf. Arrows indicate staining of collagen fibers. The boxed areas are shown at higher magnification below. (E) Concentrations of two proinflammatory cytokines, TNF-α and IL-6, in the ovary of βA mutants at 240 dpf (n = 3). (F) Expression of genes involved in apoptosis (*casp3a* and *p53*), ovarian tumors (*pawr*, *pax8*, *pgr*, *erbb2* and *cdkn2a/b*) and fibrosis (*tgfb1a* and *pparg*) in the βA mutants at 240 dpf (*p < 0.05; **p < 0.01; n = 5).

### Normal spermatogenesis and fertility in activin βA/B-deficient males

In contrast to the critical roles of activin genes in female reproduction, we found that disruption of activin β subunits, including βA (*inhbaa* and *inhbab*) and βB (*inhbb*), did not cause any obvious abnormalities in testis development and spermatogenesis. No significant differences were observed in the testes of *inhbaa* and *inhbab* single and double mutants at 120 and 240 dpf in terms of histological structure and amount of mature spermatozoa in the testis lumina determined as we reported before [45, 69] (Fig. S2A and B). The βA double mutant males (*inhbaa−/−;inhbab−/−* ♂) were also fertile when tested with double mutant females (*inhbaa−/−;inhbab−/−* ♀) although the number of eggs produced by each double mutant female was small (Fig. 1G). Similarly, the testis was also normal in the βB mutant (*inhbb−/−*) at 180 dpf in terms of structure and sperm production (Fig. S2C), and the male mutant was also fertile when tested with the female mutant (*inhbb−/−*♀ x *inhbb−/−*♂) (Fig. 1G).

### Gene expression in the hypothalamus-pituitary-gonad axis of activin mutants

Activin was first known to stimulate pituitary FSH release in mammals. To demonstrate if the loss of activin subunits has any impacts on pituitary gonadotropins in fish, we analyzed the expression of pituitary FSHβ (*fshb*) and LHβ (*lhb*) subunits in different activin β null mutants. At 90 and 120 dpf, the expression of *fshb* increased significantly in the pituitary of female βA double mutant (*inhbaa−/−;inhbab−/−*). The expression was also high in *inhbaa* single mutant (*inhbaa−/−*) albeit without statistical significance at 90 and 120 dpf; however, the increase became significant at later stages (180 and 240 dpf). In contrast, no changes were observed in *lhb* expression in either single or double βA mutant (Fig. 14A and S3A). In situ hybridization on βA double mutant (*inhbaa*−/−*;inhbab*−/−) at 90 dpf confirmed the increase of *fshb* expression (Fig. 14B and C). In contrast to that in the activin βA mutants, the expression of *fshb* and *lhb* in the activin βB mutant (*inhbb−/−*) showed no difference from the control in both males and females at 180 dpf (Fig. S5B).

**Figure 14.**
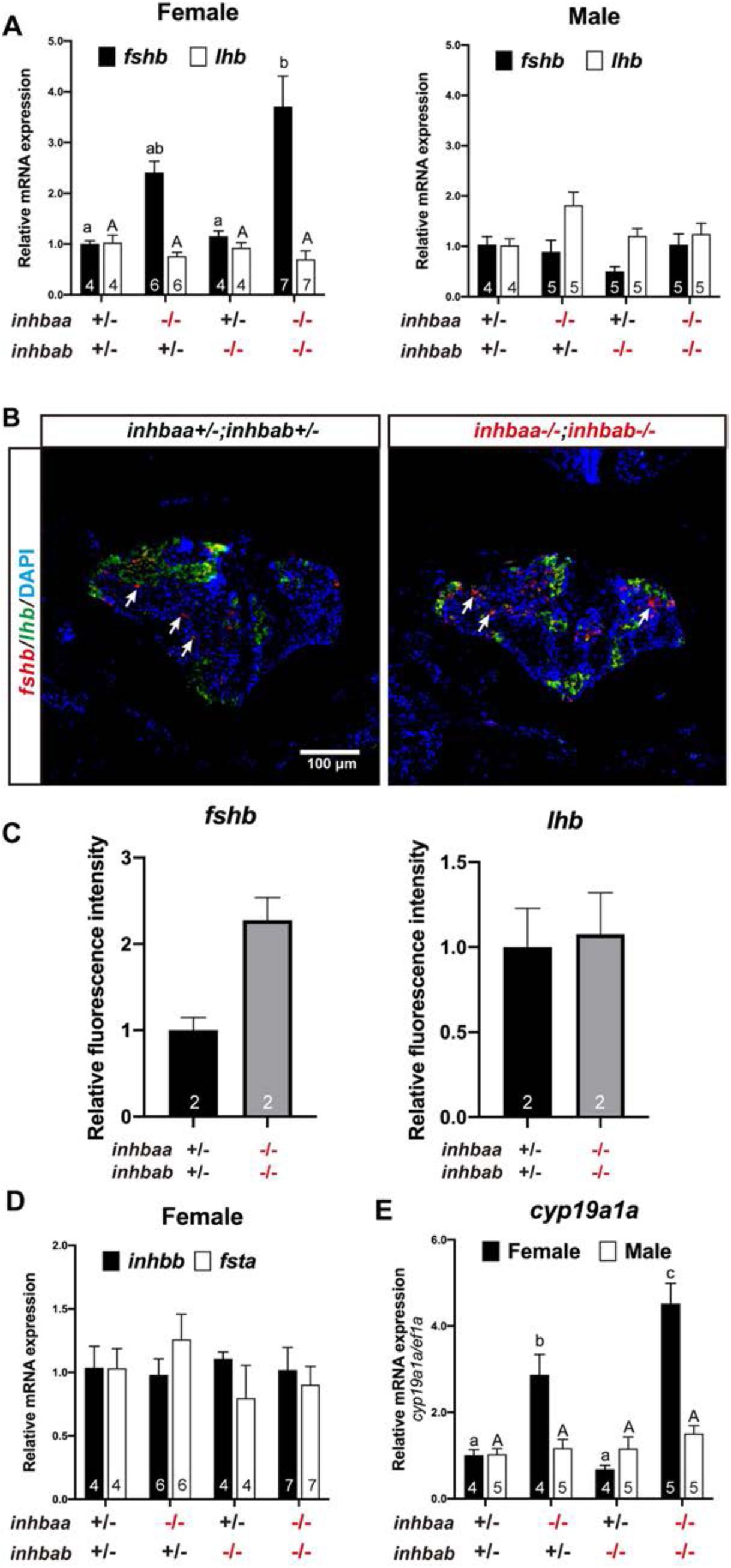
Expression analysis of pituitary and gonadal genes in activin β mutants. (A) Expression of *fshb* and *lhb* in the pituitary of activin βA mutants at 90 dpf. (B) Fluorescence in situ hybridization for *fshb* (arrows) and *lhb* in female pituitaries of the control (*inhbaa*+/−;*inhbab*+/−) and double mutant (*inhbaa*−/−;*inhbab*−/−) at 90 dpf. (C) Quantification of the in situ hybridization signals in the double mutant (n = 2). (D) Expression of *inhbb* and *fsta* in the pituitary of activin βA female mutants at 90 dpf. (E) Expression of aromatase (*cyp19a1a*) in the ovary and testis of the βA mutants at 90 dpf. Sample size is indicated in each column, and different letters in each dataset indicate statistical significance (p < 0.05).

In addition to *fshb* and *lhb*, we also examined the expression of *inhbb* and *fsta* in the pituitary at 90 dpf as they are the only two genes of the activin-inhibin-follistatin system that are expressed in the pituitary of zebrafish [35]. The result showed no change of *inhbb* and *fsta* expression in all three genotypes of activin βA mutants (*inhbaa*−/−, *inhbab*−/−, and *inhbaa*−/−*;inhbab*−/−) (Fig. 14D).

We further analyzed the expression of some essential gonadal genes in activin βA mutants (*inhbaa*−/−*, inhbab*−/− and *inhbaa*−/−*;inhbab*−/−) at 90 dpf and βB mutant (*inhbb*−/−) at 180 dpf, including FSH and LH receptors (*fshr* and *lhcgr*), estrogen β receptors (*esr1, esr2a* and *esr2b*), anti-Müllerian hormone (*amh*), doublesex and mab-3 related transcription factor 1 (*dmrt1*), androgen receptor (*ar*), and all members of the activin-inhibin-follistatin system, including *inha*, *inhbaa, inhbab*, *inhbb*, *fsta* and *fstb*. Most of these genes did not change their expression significantly in the ovary and testis of activin βA and βB mutants (Fig. S4). However, the expression of aromatase (*cyp19a1a*) increased significantly in the ovary but not testis of *inhbaa*−/− and the level increased further (> 4-fold) in the double mutant (*inhbaa*−/−*;inhbab*−/−) (Fig. 14E), similar to the pattern of *fshb* expression in these mutants (Fig. 14A).

## Discussion

The discovery of activin-inhibin-follistatin system is considered one of major breakthroughs in reproductive endocrinology in the 1980s [1]. Inhibin was identified as a major gonadal peptide hormone that exerted a negative feedback action in the pituitary to specifically suppress FSH secretion without affecting LH [70]. Activin was discovered as a side product during inhibin purification that stimulated FSH secretion [3, 4]. However, subsequent studies have shown that activin functions as the core molecule of the regulatory system whereas inhibin acts primarily as an activin antagonist that competes with activin for the same receptors [71]. In addition to their actions in the pituitary, both activin and inhibin also work in the gonads to exert paracrine and autocrine actions to regulate gonadal functions [72], and there is evidence that they may also act in the brain to control hypothalamic gonadotropin-releasing hormone (GnRH) as well [73]. Interestingly, activin was also shown in early 1990s to function as a potent mesoderm-inducing factor in embryogenesis in *Xenopus* [5, 6]; however, this view has not been supported by gene knockout studies in the mouse [7–9], leaving this a controversial issue in developmental biology.

In the present study, we carried out a comprehensive genetic study on activins in zebrafish. Using CRISPR/Cas9 method, we knocked out all three activin subunit genes (βA: *inhbaa* and *inhbab;* and βB: *inhbb*) and analyzed the phenotypes of all three single mutants (*inhbaa−/−*, *inhbab−/−* and *inhbb−/−*) as well as their combinations of double (*inhbaa−/−;inhbab−/−, inhbaa−/−;inhbb−/− inhbab−/−;inhbb−/−*) and triple (*inhbaa−/−;inhbab−/−;inhbb−/−*) knockouts. The major discoveries are summarized and discussed below.

### Roles of activins in embryonic development

One of major discoveries in developmental biology in early 1990s was the long-sought mesoderm-inducing factor (MIF), which turned out to be activin [5, 6]. A series of seminal studies in *Xenopus* have provided unequivocal evidence for potent activities of activin in inducing mesoderm formation [74, 75]. Activin as a morphogen can induce differentiation of different mesoderm tissues or organs depending on its concentration [76, 77]. However, the role of activin in embryogenesis, especially mesoderm formation, is not supported by the evidence from the mouse model. Deletion of activin subunit genes in the mouse did not lead to failure of embryonic development including mesoderm formation. The homozygous activin βA mutant was born without whiskers and lower incisors [8], and the mutant of activin βB gene was viable with some individuals showing defects in eyelid closure and female fertility [9]. Double mutant of activin βA and βB showed additive defects of two single mutants without additional abnormalities [8]. Given the conflicting results on activin involvement in embryogenesis in *Xenopus* and mice, its role in mesoderm formation needs to be validated in other species.

Our results in the present study showed that none of the three activin subunits was indispensable alone for embryonic development. However, the loss of *inhbaa* resulted in a higher mortality after 12 dpf, and double mutation of the two βA subunits (*inhbaa−/−;inhbab−/−*) further increased the death rate, suggesting an important role for activin βA in larval and post-larval development. This is similar to the report in mice that activin βA-deficient mice could develop to term but died within 24 h after birth [8]. Despite the high larval or juvenile mortality associated with *inhbaa−/−*, the overall embryonic and post-hatching development of βA mutants (*inhbaa−/−, inhbab−/−* and *inhbaa−/−;inhbab−/−*) appeared normal, indicating normal formation of mesoderm and its derived organs. Interestingly, despite being one of the most conserved regulatory proteins in vertebrates [78], the loss of activin βB (*inhbb*) did not cause any abnormalities in zebrafish growth and development, in contrast to the βA mutants. This again is similar to the loss of βB in mice, which only resulted in defects in eyelid development [9].

The dispensable roles of activin subunit genes in mesoderm formation and embryonic development was further confirmed by triple knockout of all three β subunit genes (*inhbaa−/−;inhbab−/−;inhbb−/−*), which could develop normally up to 10 dpf but not beyond 15 dpf. The mortality might primarily be due to the loss of *inhbaa* as *inhbaa−/−* was the only one among the three that showed reduced larval survival rate. The complete death of the triple mutant after 10 dpf indicates clearly compensatory functions of the three β subunits during post-hatching development. The exact causes for the death are unknown. Our data therefore support the studies in the mouse model that the lack of activin subunits did not affect mesoderm formation.

The discrepancy between the *Xenopus* and mouse models on roles of activin in mesoderm formation has prompted a view that in addition to zygotic activins, the maternal activins may contribute to the developmental process [79]. Using two activin dominant-negative variants which inhibited activin activity and depleted activin pool respectively, a study in medaka fish *Oryzias latipes* demonstrated that it was the maternally- not zygotically-derived activin that was involved in inducing mesoderm formation [80]. However, our data in zebrafish do not seem to support the involvement of maternal activins. First, no activin subunit transcripts could be detected in the oocyte of FG follicles. Second, all single (*inhbaa−/−, inhbab−/−* and *inhbb−/−*) and double mutants females (*inhbaa−/−;inhbab−/−*, *inhbaa−/−;inhbb−/−* and *inhbab−/−;inhbb−/−*) could spawn with mutant males (♀−/− x ♂−/−) to produce normal offspring. Unfortunately, we could not examine the triple mutant (*inhbaa−/−;inhbab−/−;inhbb−/−*) due to its complete mortality by sex maturation.

Having demonstrated that activins were not indispensable for embryonic development, we then focused our attention to activin involvement in reproduction. Fortunately, all single and double mutants of β subunits (*inhbaa−/−, inhbab−/−, inhbb−/−, inhbaa−/−;inhbab−/−, inhbaa−/−;inhbb−/−* and *inhbab−/−;inhbb−/−*) could survive to sexual maturity, allowing us to explore their functional importance in reproduction before and after sexual maturation.

### Local paracrine factors in ovarian follicles

Activin and inhibin were first isolated from mammalian follicular fluid [3, 4]. The expression of activin subunits has been localized primarily to the granulosa cells and sometimes the thecal cells as well [81, 82]. In fish models, activin β subunits were first localized in the somatic follicle cells of FG follicles by immunocytochemical staining in goldfish [83]. Using RT-PCR, we previously demonstrated that activin βAa (*inhbaa*) and βB (*inhbb*) subunits were both expressed exclusively in the somatic follicle cells of FG follicles whereas activin receptors and its intracellular signaling molecules Smad2/3/4/7 were abundantly expressed in the oocyte, suggesting a potential intra- follicular paracrine signaling pathway that mediates regulation of oocytes by the follicle cells [30, 31]. In this study, we showed that *inhbab*, which was not included in our previous study, was also expressed exclusively in the follicle cells together with *inhbaa* and *inhbb* as well as *inha,* further supporting the idea that activins are likely an important family of growth factors that mediate follicle cell-to-oocyte signaling. Zebrafish has two forms of follistatin (*fsta* and *fstb*), an activin-binding protein. Interestingly, *fsta* was exclusively expressed in the oocyte whereas *fstb* was equally expressed in the oocyte and follicle cells. As an important paracrine factor in the follicle, the activity of activin is likely controlled tightly by both inhibin from the follicle cells and follistatin from the oocyte (Fig. 15A).

**Figure 15.**
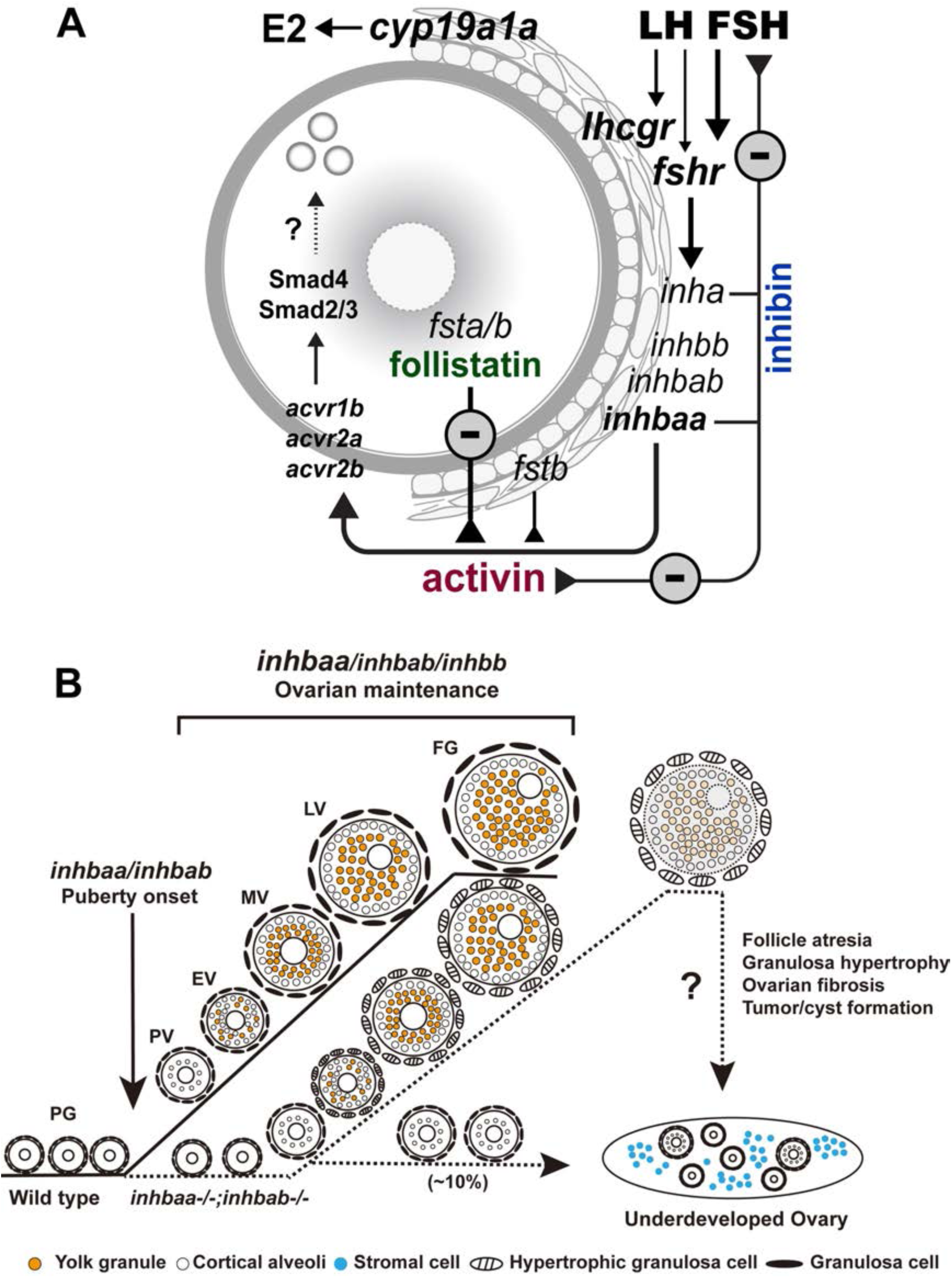
Working model on activin-inhibin-follistatin system in zebrafish ovary. (A) Intrafollicular distribution of the activin-inhibin-follistatin system. All activin/inhibin β subunits (*inhbaa*, *inhbab* and *inhbb*) are exclusively expressed in the somatic follicle cells together with inhibin α subunit (*inha*). In contrast, follistatin a (*fsta*) is exclusively expressed by the oocyte whereas *fstb* is expressed in both cell types. Activins from the follicle cells may act directly on the oocyte to control its gene expression. In addition to antagonizing activin, inhibin from the follicle cells also feeds back to the pituitary to control gonadotropin expression and secretion especially FSH. (B) Roles of activin/inhibin β subunits in folliculogenesis. Activin plays important roles in controlling follicle activation and maintaining normal folliculogenesis. Activin βA (*inhbaa* and *inhbab*) appears more important than βB (*inhbb*), especially *inhbaa* whose functions can be partially compensated by *inhbab*. The βAa (*inhbaa*) may also be the main β subunit that contributes to the formation of inhibin (αβ).

The intrafollicular distribution of activin-inhibin-follistatin in FG follicles is largely supported by a recent single-cell transcriptome study on early follicles (40 dpf), which revealed finer distribution of the activin subunits at single cell level before follicle activation. The *inhbaa-*expressing cells were the most abundant among all activin-inhibin subunits and its expression was primarily concentrated in the granulosa cells with some signals also detectable in the theca and stromal cells; however, *inhbab*- expressing cells were much less abundant, and its expression was almost exclusively detected in the theca cells. By comparison, *inhbb* was mostly expressed in the stromal and germ cells (oocytes) with weak signals in the granulosa and theca cells. Interestingly, the expression of *inha* was exclusively localized in the granulosa cells, coexisting mostly with that of *inhbaa* [84]. These data suggest that in zebrafish ovary, at least in newly-formed young follicles at 40 dpf, different forms of activin molecules (activin A, B and AB) can be formed in all types of cells including granulosa, theca and stromal cells as well as the oocytes. However, inhibin may be primarily produced and secreted by the granulosa cells as inhibin A because of the coexistence of *inha* and *inhbaa* expression in the granulosa cells. This view is further supported by our observation that *inha* and *inhbaa* mutants shared some phenotypes at both pituitary and ovary levels (see discussions below). The expression of *inhbb* in the germ cells at 40 dpf but not in the FG oocytes of mature fish (> 90 dpf) suggests that the spatial expression pattern of activin subunits may change during ovarian development and folliculogenesis.

### Involvement in follicle activation and puberty onset in females

We have previously shown that during follicle activation or PG-PV transition, activin subunits, especially βAa (*inhbaa*), showed a significant increase in expression [32, 33], suggesting a potential role for activins in follicle activation or puberty onset, which is marked by appearance and accumulation of cortical alveoli in the leading follicles [56]. Surprisingly, none of the three single mutants of activin subunits showed any changes in the timing of follicle activation, which normally occurs around 45 dpf when the body size reaches 1.8 cm in BL and/or 100 mg in BW [56, 57]. Interestingly, many individuals of the βA double mutant (*inhbaa*−/−;*inhbab*−/−) showed a significant delay in follicle activation as the PV follicles were often absent in individuals with body size far beyond the threshold for triggering puberty onset (*e.g*., 2.2 cm/138 mg). Furthermore, in some double mutant females, the follicles were completely blocked at PG or PV stage. This occurred in about 10% mutant females examined, suggesting incomplete penetrance. The abnormalities in βA double but not single mutants suggest compensatory functions of the two βA subunits (*inhbaa* and *inhbab*) in promoting follicle activation and subsequent PV follicle development. Since the triple mutant of activin β subunits (*inhbaa*−/−;*inhbab*−/−;*inhbb*−/−) showed post-larval lethality, we were not able to assess the impact of complete lack of activin subunits on follicle activation.

The view that activins may promote follicle activation agrees well with our recent report that the loss of inhibin (*inha−/−*) advanced follicle activation significantly. Interestingly, the mutation of *inha* caused a nearly 5-fold increase in *inhbaa* expression in the ovary, suggesting an increased production of activin A [45]. We therefore hypothesize that activin and inhibin are at a dynamic equilibrium during follicle activation, and the loss of either one will disrupt the balance of the two antagonistic regulators, leading to delayed or advanced (precocious) puberty onset, respectively (Fig. 15).

### Long-term maintenance of ovarian structure and function

Activin subunits especially *inhbaa* exhibit dynamic changes in expression during folliculogenesis in zebrafish. The expression of *inhbaa* starts to increase during PG-PV transition and continues to rise during follicle growth [32], suggesting roles for activins not only at follicle activation as described above, but also during follicle growth. Surprisingly, examination of post-pubertal ovaries showed no abnormalities in follicle development in any single mutants (*inhbaa*−/−, *inhbab*−/− and *inhbb*−/−) up to sexual maturation. However, in βA double mutant (*inhbaa*−/−*;inhbab*−/−), we often observed hypertrophy of granulosa cells and hyperplasia of inter-follicular stromal cells. Interestingly, the granulosa cell hypertrophy somehow phenocopied inhibin mutant (*inha−/−*) [45]. This may be because the mutation of βA might also cause loss of inhibin A (αβA), resulting in partial phenocopying of the *inha−/−* mutant. As discussed above, *inhbaa* may be the major form of β subunits that contribute to the biosynthesis of inhibin in zebrafish. The single cell transcriptome data showing coexistence of *inha* and *inhbaa*, but not *inhbab* and *inhbb*, in the granulosa cells further support this idea [84].

Despite the abnormalities in βA null fish, the double mutant females were fertile at 120 dpf. However, the condition of the ovary deteriorated quickly after 180 dpf in both single (*inhbaa*−/−) and double (*inhbaa*−/−*;inhbab*−/−) mutants, showing follicle degeneration or atresia, stromal cell hyperplasia and reduction in fecundity. The strong TUNEL signals in the stromal cells and increased expression of Caspase-3 mRNA (*casp3a*) and protein in *inhbaa*−/− ovary suggest an important role for activin βA in ovarian tissue homeostasis and long-term maintenance of ovarian structure and function. This is strongly supported by the evidence that the ovaries of *inhbaa*−/− fish regressed significantly with folliculogenesis arrested at PG stage at 180 mpf (> 540 dpf). The ovarian regression and disorganization were also accompanied with increased fibrosis and inflammation. In mammals, activin is associated with apoptotic activities in a wide range of cell types and tissues [85, 86] and it is also widely involved in tissue repair, fibrosis and inflammation [87, 88]. The increased expression of inflammatory cytokines (TNF-α and IL-6) and genes involved in fibrosis such as *tgfb1a* in *inhbaa*−/− ovary supports a conservative role for activin in tissue homeostasis.

Similar to activin βA, the βB subunit (*inhbb*) also played a role in ovarian and follicular maintenance. However, the impact of its loss on the ovary and female fertility became significant at much later stage than that of βA mutants. Since *inhbb* expression showed little overlapping with that of *inha* in early follicles [84], the loss of *inhbb* might not affect inhibin biosynthesis, in contrast to that of *inhbaa*. The mild phenotype of βB mutant as compared to the βA mutants agrees well with the situation in mice. In contrast to the βA-deficient mice, which died within 24 h after birth, the βB null mice could develop to sex maturity and the females showed no abnormalities in the ovary and folliculogenesis [9].

### Potential roles of activins in ovarian disorders

As important regulators in reproduction, activin-inhibin-follistatin system has been associated with some ovarian disorders in humans. For example, human polycystic ovarian syndrome (PCOS) is associated with low serum concentration of activin and high concentration of follistatin [89]. In mice, inhibin α (INHA) was reported to function as a negative regulator of stromal cell proliferation, and its loss resulted in formation of gonadal tumors at very early stage accompanied by overexpression of activin βA subunit [90]. In zebrafish, our recent study also demonstrated tumorigenesis in the gonads of some *inha* null mutant [45]. The mechanism by which Inha acts as a tumor suppressor is unknown. As a secreted protein, Inha most likely works as inhibin. Interestingly, the *inhbaa−/−* ovaries also developed tumor- or cyst-like structures after about 12 mpf. Due to the high mortality, we did not have chance to observe formation of tumor-like tissues in the double mutant (*inhbaa*−/−*;inhbab*−/−) at the same age. Similar to inhibin, activin has also been reported to act as a potent inhibitor to suppress proliferation of human epithelial ovarian cancer cells [91, 92]. The changed expression of inhibin/activin subunits has been observed in the primary ovarian cancers of epithelial origin [93]. This, together with our results, suggests that activin may also act as a negative regulator in ovarian cell proliferation and tumorigenesis. In addition to inducing formation of ovarian cysts or tumor-like tissues, the lack of *inhbaa* also resulted in extensive fibrosis in the zebrafish ovary. This agrees well with studies in mammals. In humans, activin is involved in the pathogenesis of inflammatory and fibrotic diseases [88, 94]. Interestingly, TGF-β1a (*tgfb1a*), which was highly expressed in the *inhbaa−/−* ovary, plays a critical role in tissue fibrosis in many organs including ovary [66, 95].

### Normal spermatogenesis without activins

Although activin and inhibin subunits are expressed in fish testis [27, 55, 96], none of the single mutants of activin subunits (*inhbaa*−/−*, inhbab*−/− and *inhbb−/−*) showed phenotypic abnormalities in males in terms of testis development, maintenance and spermatogenesis. The spermatogenesis was normal in mutant males, which were functionally fertile. Despite the high lethality of activin βA double mutants (*inhbaa*−/−*;inhbab*−/−), a small number of the mutant fish could survive to sexual maturity. Like the single mutants, the βA double mutant males were also fertile with normal spermatogenesis in the testis. In mice, conditional knockout of activin A in fetal Leydig cells resulted in decreased Sertoli cell proliferation and failed cord elongation and expansion in the fetal testis [26].

### Regulation of gene expression in the hypothalamic-pituitary-gonadal (HPG) axis

Activin was originally discovered in the ovarian follicular fluid as a specific stimulator of pituitary FSH without any effect on LH, in contrast to inhibin that suppresses FSH secretion [3, 4]. Subsequent studies have shown that both activin and inhibin have widespread actions at all levels of the HPG axis to regulate expression of a variety of genes. In the hypothalamus, activin stimulates GnRH expression and release from both hypothalamic explants [97] and GnRH-secreting neuronal cell lines [98]. In the pituitary, in addition to stimulating FSH biosynthesis and secretion, activin also increases transcription of GnRH receptors in gonadotrophic cells [99] and therefore potentiates GnRH-stimulated FSHβ expression [100]. In the ovary, activin acts as a multifunctional factor regulating various aspects of ovarian functions including oogonial proliferation and primordial follicle formation, follicle growth and development, oocyte maturation and ovulation, and steroidogenesis [82]. In this study, we also examined expression changes of some genes that are critical to reproduction in the pituitary and gonads in various activin β mutants.

We first looked at the response of pituitary gonadotropins especially FSH (*fshb*) to the loss of activin subunits as this is the primary function for which activin was discovered. In agreement with that in mammals, activin also stimulates *fshb* expression in fish including zebrafish *in vitro* [27, 35, 36]. To our surprise, instead of decrease as expected, the expression of *fshb* in βAa single (*inhbaa*−/−) and double mutant (*inhbaa*−/−*;inhbab*−/−) increased significantly compared to the control and βAb single mutant (*inhbab*−/−). This again phenocopied the *inha* mutant (*inha*−/−) which also showed an increased *fshb* expression [45]. Since the loss of β subunits especially *inhbaa* may also result in the loss or reduction of inhibin, it is conceivable that both *inhbaa* and *inha* mutants increased *fshb* expression. In contrast to βA mutants, the βB mutant (*inhbb*−/−) showed no changes in *fshb* expression despite its potential role in the pituitary in mediating inhibin and follistatin regulation of *fshb* expression as a local paracrine factor [35].

Also surprisingly, the expression of LH (*lhb*) remained unchanged in all activin mutants although our previous studies have consistently shown opposite effects of activin on the expression of *fshb* and *lhb* in both zebrafish [35] and goldfish [36], *i.e.,* stimulation of *fshb* but suppression of *lhb*. This has also been confirmed in other fish species such as European eel [37]. Our results in activin mutants indeed agree well with that in mammals. In mice, loss of activin β subunits also increased the concentration of FSH in serum with no change in LH concentration [9, 23]. The lack of *lhb* response to the loss of activin subunits in zebrafish suggests that the regulation of *lhb* by activin could have been compensated by other regulatory mechanisms in the mutants. Alternatively, the inhibition of *lhb* expression by activin in vitro may represent an acute and short-term response of LH to activin exposure.

We have proposed that there exists a local activin-follistatin system in goldfish and zebrafish pituitaries that regulates FSH and LH biosynthesis [35, 101, 102]. In zebrafish, it is βB *inhbb* subunit that is expressed in the pituitary, forming a local paracrine mechanism controlling gonadotropin biosynthesis [35], similar to the situation in the rat [103, 104]. A similar mechanism has also been reported in mammals [105] and other fish species such as the grass carp [106]. The lack of response of *fshb* and *lhb* to *inhbb* mutation suggests that such paracrine mechanism may work on short-term basis during zebrafish daily reproductive cycle, and the long-term loss of *inhbb* in the mutant might have been compensated by other regulatory mechanisms in vivo.

We also examined the expression of a variety of genes in the gonads of activin mutants, including gonadotropin receptors (*fshr* and *lhcgr*), estrogen receptors (*esr1*, *esr2a* and *esr2b*), and all members of the activin-inhibin-follistatin system (*inhbaa*, *inhbab*, *inhbb*, *inha*, *fsta* and *fstb*). Most of these genes remained unchanged or changed slightly in the mutant. However, the expression of estrogen-producing enzyme aromatase (*cyp19a1a*) showed a significant increase in both βAa single (*inhbaa*−/−) and βA double mutant (*inhbaa*−/−*;inhbab*−/−) ovaries, which again phenocopied the inhibin mutant (*inha−/−*) [45]. Whether the increased expression of *cyp19a1a* was due to the increased production of FSH in the pituitary or direct actions of activin in the ovary remains unknown.

In summary, we successfully knocked out all the activin/inhibin β subunit genes in the zebrafish using CRISPR/Cas9 technology. Our results showed normal embryonic development in all single, double and triple mutants. The loss of activin βA or βB had no effect on spermatogenesis and testis development in males. However, mutation of βA showed profound effects on female reproduction, especially *inhbaa−/−*. The expression of *fshb* in the pituitary and *cyp19a1a* in the ovary increased significantly in activin βA mutant. Despite this, the βA double mutant (*inhbaa−/−;inhbab−/−*) showed delayed puberty onset, somatic cell hyperplasia and granulosa cell hypertrophy. Other ovarian disorders also developed in βA mutants at later stage of life including follicle degeneration or atresia, ovarian fibrosis, cyst/tumor formation, and cessation of folliculogenesis. In contrast to the βA mutants, the βB mutant (*inhbb−/−*) showed much milder phenotypes with reduced female fertility only at later stage. Our comprehensive genetic data provided substantial information about the functions of the inhibin-activin- follistatin system in fish development and reproduction.

## Acknowledgements

We thank Ms. Phoenix Un Ian LEI for the maintenance and management of the zebrafish facility and the Histology Core of the Faculty of Health Sciences for technical support. This study was supported by grants from the University of Macau (MYRG2019-00123-FHS, MYRG2020-00192-FHS and CPG2020-00005-FHS) and The Macau Fund for Development of Science and Technology (FDCT173/2017/A3 and FDCT0132/2019/A3) to WG. KW is supported by the Macau Young Scholars Program (AM2020025).

---

**Figure S1.**
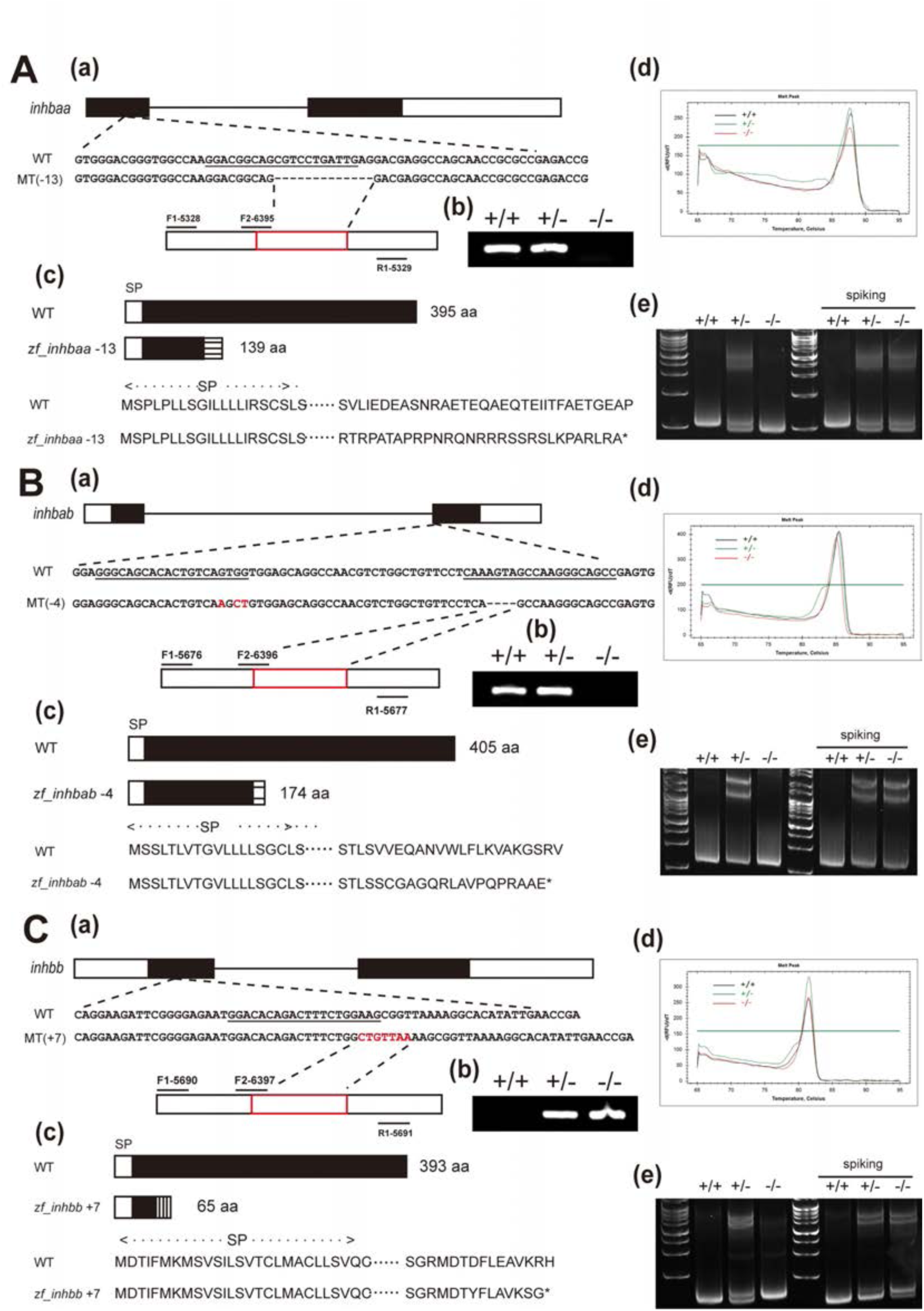
Mutagenesis of zebrafish *inhbaa*, *inhbab* and *inhbb.* (A) Mutagenesis of *inhbaa* and mutant characterization. (a) Schematic illustration of the genomic structure of zebrafish *inhbaa* gene. The underlined sequence indicates CRISPR/Cas9 target site. The dashed line indicates the deleted sequence (−13 bp) of zebrafish *inhbaa*. (b) The expression of mutated transcript in the ovary. RT-PCR was performed on total RNA extracted from the ovary with a mutant-specific primer (F2-6395) overlapping with the deleted sequence. (c) Schematic representation of *inhbaa* amino acid sequence. The mutation is expected to introduce a premature stop codon (*). (d) Genotyping by HRMA with the primer pairs of F1-5328 and R1-5329; (e) HMA confirmation of different genotypes of *inhbaa* mutant. (B) Mutagenesis of *inhbab* and mutant characterization. (a) Schematic illustration of the genomic structure of zebrafish *inhbab* gene. The dashed line indicates the deleted sequence (−4 bp). (b) RT-PCR confirmation of mutation at the transcript level with a mutant-specific primer (F2-6396). (c) Schematic representation of *inhbab* amino acid sequence. (d) Genotyping by HRMA with the primer pairs of F1-5676 and R1-5677; (e) HMA confirmation of different genotypes of *inhbaa* mutant. (C) Mutagenesis of *inhbb* and mutant characterization. (a) Schematic illustration of the genomic structure of zebrafish *inhbb* gene. The inserted nucleotides are marked in red (+7 bp). (b) RT-PCR confirmation of mutation at the transcript level with a mutant-specific primer (F2-6397). (c) Schematic representation of *inhbb* amino acid sequence. (d) Genotyping by HRMA with the primer pairs of F1- 5690 and R1-5691; (e) HMA confirmation of different genotypes of *inhbb* mutant. Since the homozygous mutant (−/−) and WT (+/−) were sometimes difficult to distinguish by HRMA, we spiked the samples with WT DNA to generate heterozygous product in mutant samples before PCR amplification. WT, wild type; MT, mutant.

**Figure S2.**
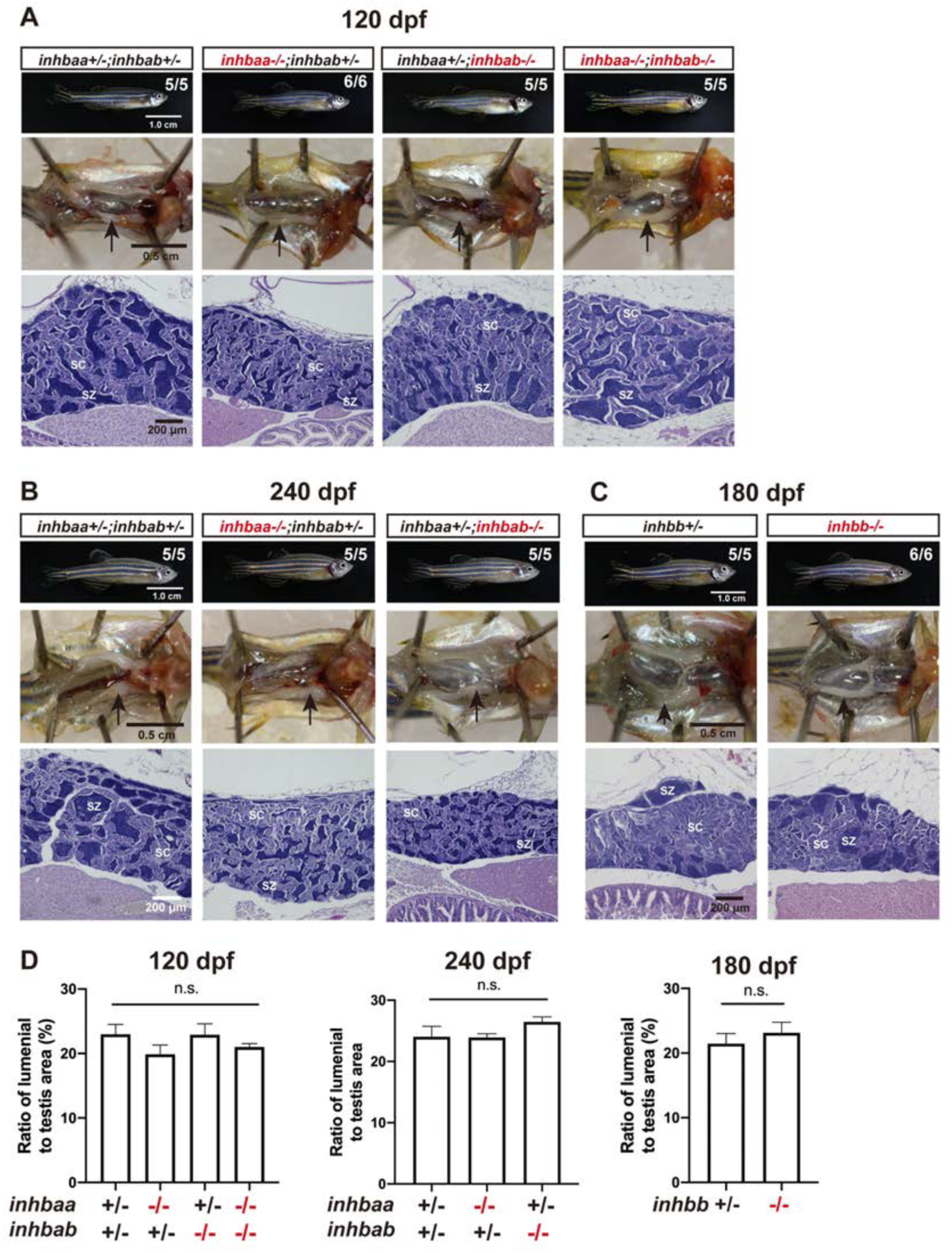
Normal spermatogenesis in activin βA mutant males. (A) Morphology, gross anatomy, and histological structure of activin βA mutant males (*inhbaa−/−, inhbab−/−* and *inhbaa−/−;inhbab−/−*) at 120 dpf. (B) Activin βA single mutant males(*inhbaa−/−, inhbab−/−*) at 240 dpf. (C) Activin βB mutant males (*inhbb−/−*) at 180 dpf. (D) Quantification of spermatozoa-filled luminal spaces in the testis at 120 and 240 dpf for activin βA mutant males and 180 dpf for βB mutant males (n = 4).

**Figure S3.**
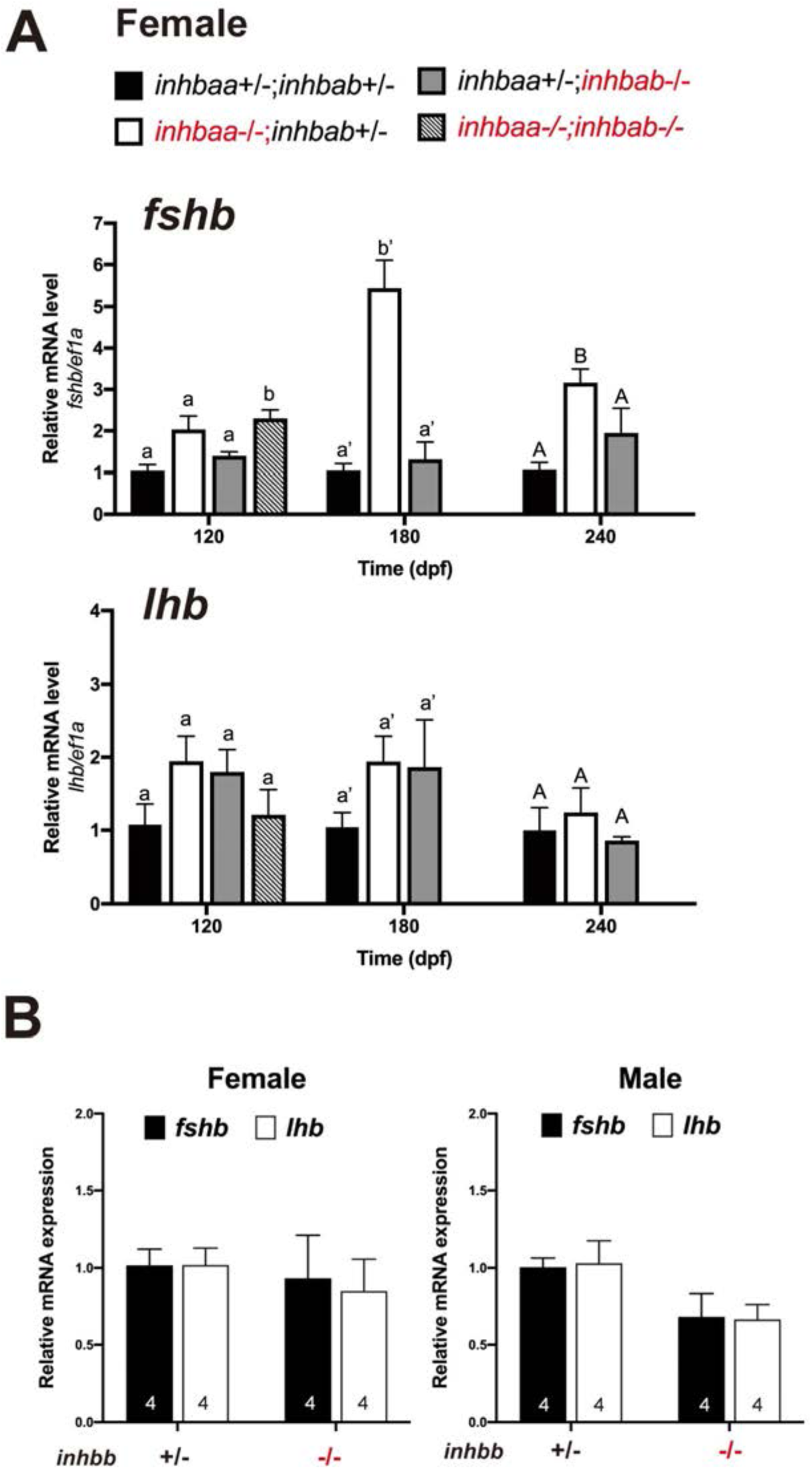
Expression of gonadotropins (*fshb* and *lhb*) in the pituitary of activin β mutants. (A) Expression of *fshb* and *lhb* in the pituitary of female βA mutants at 120, 180 and 240 dpf (n=3). The double mutant (*inhbaa−/−;inhbab−/−*) was not included for 180 and 240 dpf due to high mortality. Different letters in each dataset indicate statistical significance (p < 0.05). (B) Expression of *fshb* and *lhb* in the pituitary of male and female βA (*inhbb*) mutant at 180 dpf.

**Figure S4.**
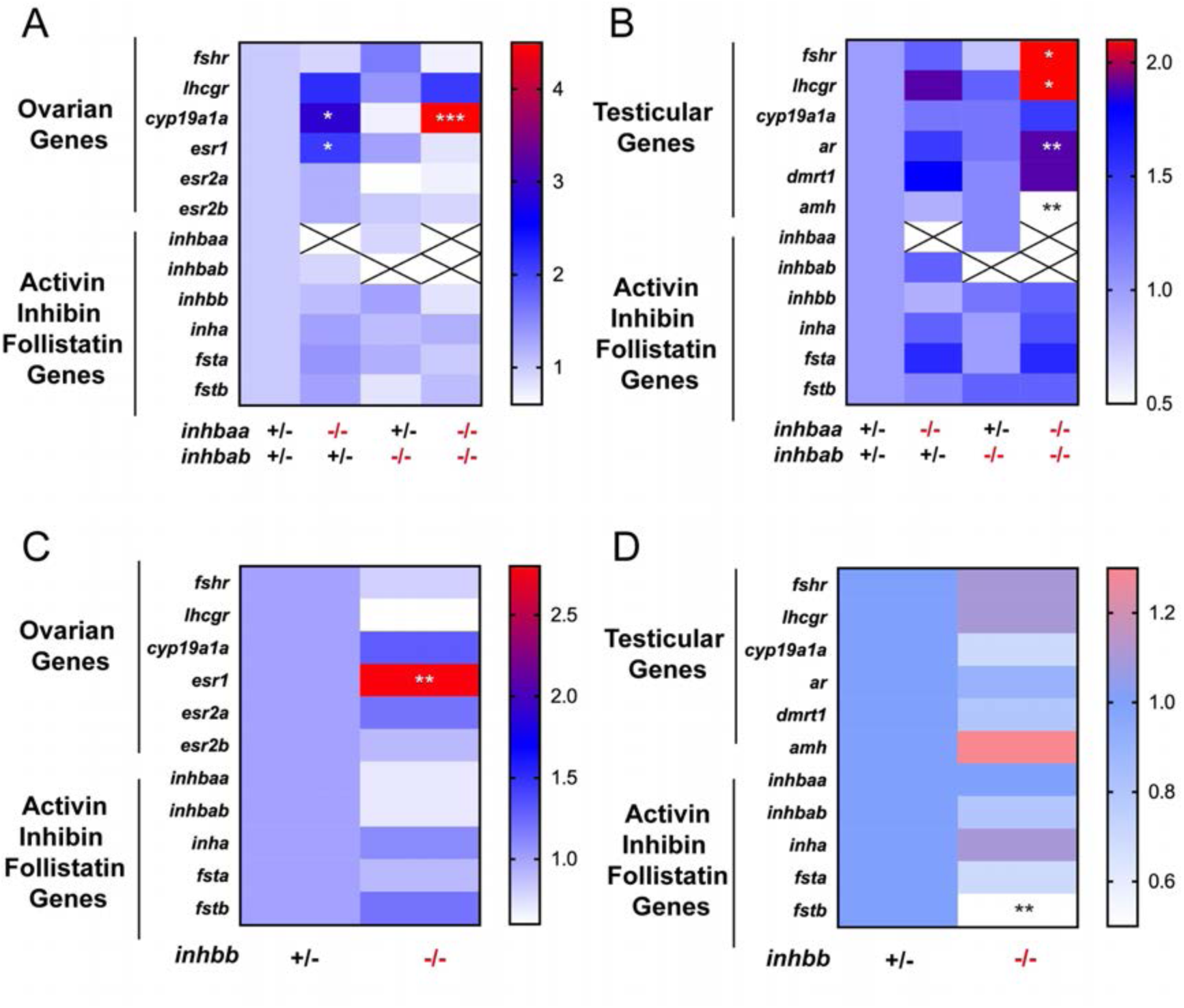
Expression of selected gonadal genes and members of the activin-inhibin-follistatin system in the ovary and testis of activin βA and βB mutants. (A) Gene expression in the ovary of activin βA (*inhbaa, inhbab*) mutants at 90 dpf (n = 4-5). (B) Gene expression in the testis of the βA mutants at 90 dpf (n = 5). (C) Gene expression in the ovary of βB mutant (*inhbb*) at 180 dpf (n = 4). (D) Gene expression in the testis of the βB mutant at 180 dpf (n = 4). The expression levels are normalized to *ef1a* and presented as the fold change compared with the control fish. The asterisks indicate statistical significance (*p < 0.05, **p < 0.01 and ***p < 0.001).

**Table S1.**
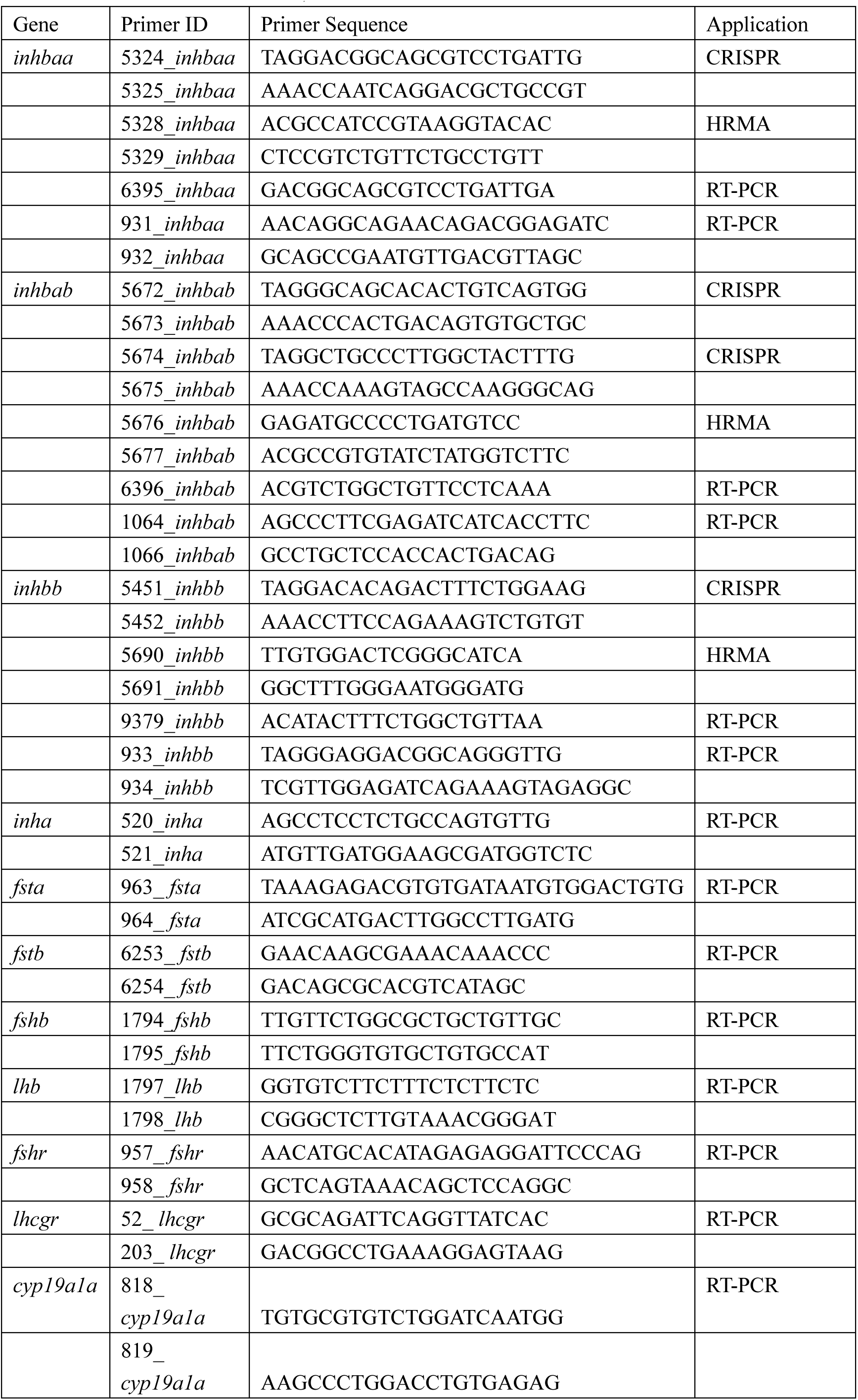
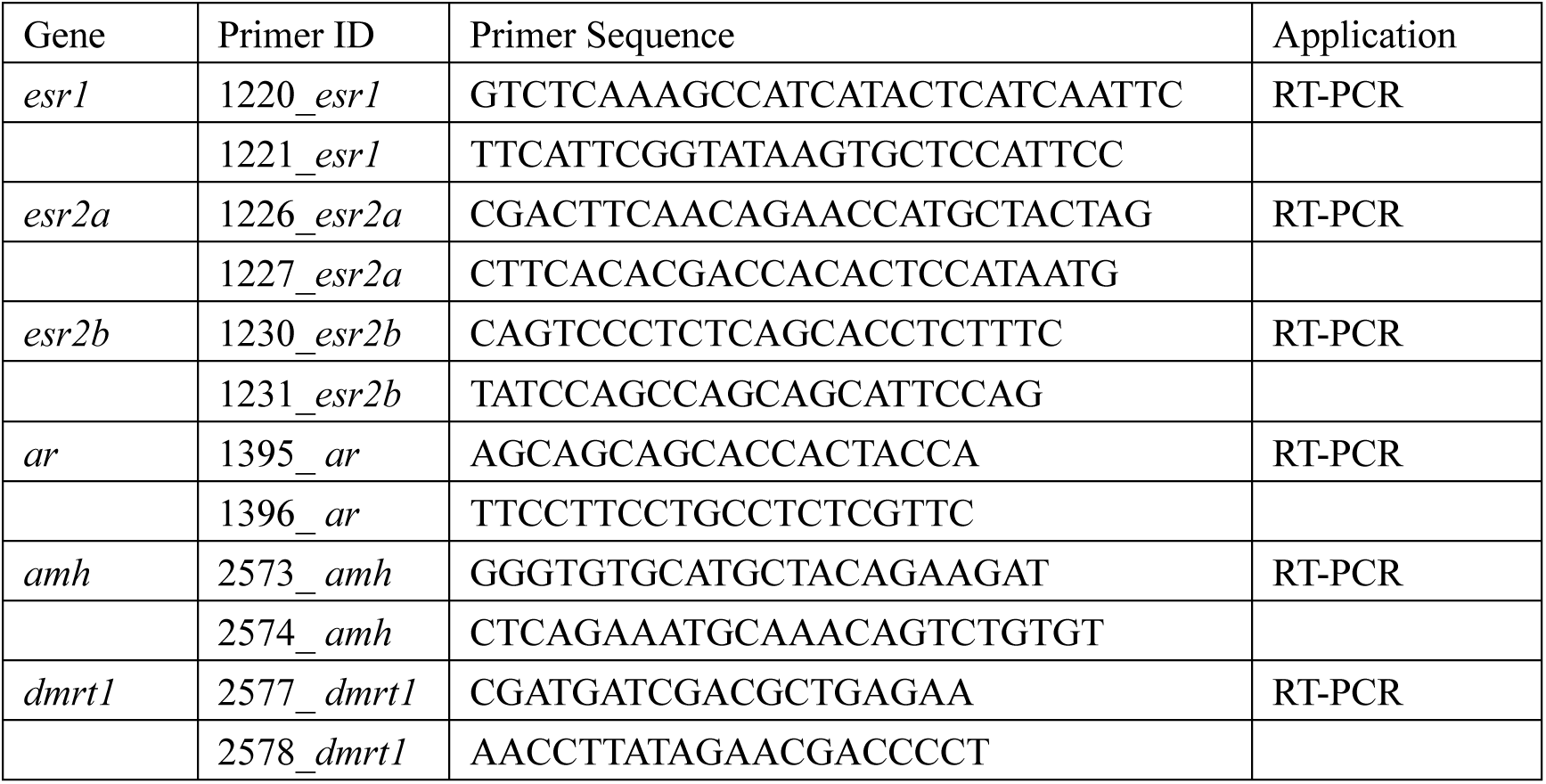
Primer used for CRISPR, HRMA and RT-PCR

